# Does Sex Modify an Association of Electrophysiological Substrate with Sudden Cardiac Death? The Atherosclerosis Risk in Communities (ARIC) Study

**DOI:** 10.1101/674689

**Authors:** Stacey J. Howell, David German, Aron Bender, Francis Phan, Srini V. Mukundan, Erick A. Perez-Alday, Nichole M. Rogovoy, Kazi Haq, Katherine Yang, Ashley Wirth, Kelly Jensen, Larisa G. Tereshchenko

## Abstract

**Background:** Sex is a well-recognized risk factor for sudden cardiac death (SCD). Sex differences in electrophysiological (EP) substrate of SCD are known. However, it remains unknown whether sex can modify an association of EP substrate with SCD.

**Methods:** Participants from the Atherosclerosis Risk in Communities study with analyzable ECGs (n=14,725; age, 54.2±5.8 yrs; 55% female, 74% white) were included. EP substrate was characterized by traditional 12-lead ECG (heart rate, QRS, QTc, Cornell voltage), spatial ventricular gradient (SVG) and sum absolute QRST integral (SAI QRST) metrics. Two competing outcomes were adjudicated SCD and nonSCD. Interaction of ECG metrics with sex was studied in Cox proportional hazards and Fine-Gray competing risk models. Relative hazard ratio (RHR) and relative sub-hazard ratio (RSHR) with a 95% confidence interval for SCD and nonSCD risk for women relative to men were calculated. Model 1 was adjusted for prevalent cardiovascular disease (CVD) and risk factors. Time-updated model 2 was additionally adjusted for incident non-fatal CVD.

**Results:** Over a median follow-up of 24.4 years, there were 530 SCDs (incidence 1.72 (1.58-1.88)/1000 person-years) and 2,178 nonSCDs (incidence 7.09; (6.80-7.39)/ 1000 person-years). Women experienced a greater than men risk of SCD associated with Cornell voltage (RHR 1.18(1.06-1.32); P=0.003), SAI QRST (RHR 1.16(1.04-1.30); P=0.007), area SVG magnitude (RHR 1.24(1.05-1.45); P=0.009), and peak SVG magnitude (RHR 1.22(1.04-1.44); P=0.018), independently from incident CVD. Greater risk of SCD for women than men associated with QRS duration (RHR 1.24(1.07-1.44); P=0.004) and QTc (RSHR 1.15(1.02-1.30); P=0.025) was explained by incident CVD. Furthermore, women had greater odds of SCD associated with heart rate (RSHR 1.19(1.01-1.40); P=0.036), independently of incident CVD.

**Conclusions:** Sex modifies an association of EP substrate with SCD. In women, global EP substrate is associated with up to 27% greater risk of SCD than in men. Development of sex-specific risk scores of SCD is necessary. Further studies of mechanisms behind sex differences in EP substrate of SCD are warranted.

## Introduction

Sudden cardiac death (SCD) is a leading cause of death in the United States. Sex is a well-recognized risk factor for SCD.^1^ SCD more commonly occurs in men as compared to women. Women have a lower prevalence of obstructive coronary heart disease (CHD) and systolic dysfunction preceding SCD.^2^ Women are also less likely than men to receive implantable cardioverter defibrillators (ICD) for primary and secondary prevention of SCD.^3^ Women were underrepresented in ICD trials, and, in result, randomized controlled trials (RCTs) did not have sufficient statistical power to detect a significant survival benefit of ICD therapy in women.^4^ Moreover, regardless of underlying etiology of heart disease,^5^ women with ICDs are less likely to experience ventricular tachyarrhythmias,^5^ and receive appropriate ICD therapies,^6^ and are more likely to suffer device-related complications.^7^ Therefore, SCD risk stratification is especially important for women.

Risk stratification of SCD for both sexes is inadequate and current practice relies on the degree of left ventricular (LV) dysfunction.^8^ While sex differences in cardiac electrophysiology have been recognized,^9^ sex is not routinely considered a potential effect modifier of the association between electrophysiological (EP) substrate and SCD. As women develop CHD approximately 10 years later than men, women are commonly viewed as “younger men”.

Widely available routine surface 12-lead electrocardiogram (ECG) describes global characteristics of the EP substrate of SCD.^10^ Sex differences in EP substrate are known: women have faster heart rate, narrower QRS and longer QT interval than men.^9^ However, it remains unknown whether sex can modify an association of EP substrate with SCD. Recently, we expanded the armamentarium for global ECG measures of EP substrate by adding global electrical heterogeneity (GEH).^1^ GEH is quantified by spatial ventricular gradient (SVG) magnitude and direction (elevation and azimuth), its scalar value sum absolute QRST integral (SAI QRST), and spatial QRS-T angle. The addition of GEH to demographic (age, sex, race) and clinical (diabetes, hypertension, CHD, stroke) risk factors improves reclassification of SCD^1^. However, it remains unknown whether GEH as a measure of independent EP substrate is different in men and women, and whether sex can modify the association of GEH and traditional global ECG measures with SCD. We hypothesized that (1) there are sex differences in GEH, and (2) sex modifies the association of traditional and novel global ECG measures of EP substrate with SCD.

## Methods

### Study population

The Atherosclerosis Risk in Communities (ARIC) study is a prospective cohort that recruited 15,792 men and women, age 45-64 years, selected as a probability sample from four United States communities. Participants were recruited in 1987-1989. Standardized examinations were conducted as previously described.^11^ Included in the analysis were ARIC cohort participants with recorded resting 12-lead ECG and measured global electrical heterogeneity (GEH); n=15,777. Excluded were participants self-identifying as non-white or non-black race (n=48), or as black at the Washington County, and Minneapolis field centers (n=55), those with missing covariates (n=903), and non-sinus median beat (n=46). The final sample of participants with normal sinus median beat included 14,725 participants.

### Exposures of sex and electrocardiographic global electrical heterogeneity

Resting 12-lead ECGs of the first five study visits were analyzed. Visit 1 was conducted in 1987-1989, visit 2 in 1990-1992, visit 3 in 1993-1995, visit 4 in 1996-1998, and visit 5 in 2011-2013. Traditional ECG amplitudes and intervals were measured by the 12 SL algorithm (GE Marquette Electronics, Milwaukee, WI). Sex-specific Cornell product was calculated to define ECG-left ventricular hypertrophy (LVH).

GEH was measured as previously described,^12^ by spatial QRS-T angle, SVG magnitude, azimuth, and elevation, and SAI QRST. The MATLAB (MathWorks, Natick, MA, USA) software code for GEH measurement is provided at https://physionet.org/physiotools/geh. Both area and peak SVG vectors^12^ and QRS-T angles were included in analysis. Previously reported^1^ area-based GEH metrics were used in this study. To measure peak-vector-based GEH metrics, we constructed a time-coherent median beat and defined isoelectric heart vector origin point, as described.^13^ The MATLAB (MathWorks, Natick, MA, USA) software code for the heart vector origin definition is provided at https://github.com/Tereshchenkolab/Origin. In this study, we included only participants with a normal sinus median beat.

### Primary outcome: sudden cardiac death

Follow-up of ARIC participants^14^ and adjudication of SCD was previously described.^15^ Physician-adjudicated SCD was defined as a sudden pulseless condition in a previously stable individual without evidence of a non-cardiac cause of cardiac arrest if the cardiac arrest occurred out of the hospital or in the emergency room. Definite, probable, or possible SCD was included in this study as a primary outcome.

### Competing mortality outcome: non-sudden cardiac death

Non-sudden cardiac death (nonSCD) was defined as a composite of fatal CHD, heart failure (HF) death, death in a participant with baseline HF, or incident hospitalized HF. Cases of fatal CHD were adjudicated by the ARIC Morbidity and Mortality Classification Committee.^14, 16^ Baseline prevalent HF was defined as a symptomatic HF (stage 3 by the Gothenburg criteria, requiring manifestation of HF cardiac and pulmonary symptoms in addition to medical treatment^17^), or self-reported use of HF medication. Incident HF was defined based on the HF codes in a death certificate or an International Classification of Diseases (*ICD-*9) discharge code, in any position, as previously described.^18^ All other deaths comprised the noncardiac death outcome.

### Baseline clinical characteristics

Body mass index (BMI) was categorized as underweight (<18.5 kg/m^2^), normal weight (18.5 to <25.0 kg/m^2^), overweight (25.0 to <30.0 kg/m^2^) or obese (≥30.0 kg/m^2^). Hypertension was defined as blood pressure (BP) of ≥140/90 mm Hg, or report of taking antihypertensive medication at visit 1. Diabetes was defined as nonfasting blood glucose ≥200 mg/dL, fasting blood glucose ≥126 mg/dL, self-reported physician diagnosis of diabetes, or reporting taking medication for diabetes or high blood sugar at visit 1. Stages of chronic kidney disease (CKD) were based on estimated glomerular filtration rate (eGFR) calculated using the CKD Epidemiology Collaboration equation (CKD-EPI).^19^ Stage 1 CKD included participants with normal or increased kidney function (eGFRCKD-EPI ≥90 mL/min/1.73 m^2^). Stage 2 CKD included mild decreased kidney function (eGFRCKD-EPI 60 to <90 mL/min/1.73 m^2^). Stage 3 CKD included moderate decreased kidney function (eGFRCKD-EPI 30 to <60 mL/min/1.73 m^2^). Stage 4 CKD participants had severe decreased kidney function (eGFRCKD-EPI 15 to <30 mL/min/1.73 m^2^), and stage 5 CKD was established kidney failure (eGFRCKD-EPI <15 mL/min/1.73 m^2^). Physical activity during leisure time was assessed using the Baecke questionnaire.^20^ Postmenopausal status was determined by questionnaire^21^ and was defined as either surgical or natural postmenopause, or primary amenorrhea. Prevalent stroke was diagnosed by a stroke and transient ischemic attack diagnostic algorithm, as previously reported^22^. Prevalent CHD was defined as a self-reported physician diagnosis of myocardial infarction (MI), or baseline ECG evidence of MI by the Minnesota code, a history of angina pectoris, or a history of coronary revascularization (either via coronary artery bypass surgery or percutaneous coronary intervention). Use of antiarrhythmic drugs included self-reported and validated by medications inventory use of class I, II (beta-blockers), III, IV (phenylalkylamines and benzothiazepines calcium channel blockers), or V (digoxin) antiarrhythmic agents.

### Incident non-fatal cardiovascular events

Incident atrial fibrillation (AF) was defined as either detected on follow-up 12-lead ECG or hospital discharge records (*ICD-9* code 427.3).^23^ Incident stroke was physician-adjudicated, as previously described.^24^ Definite or probable incident strokes are included in this study. Expert-adjudicated incident CHD was defined as a definite or probable MI, angina, or a coronary revascularization procedure.^14, 16^ Incident HF was defined above.^18^

### Statistical analyses

#### Cross-sectional analyses at the baseline

Normally distributed continuous variables were compared using a *t*-test and presented as means ± standard deviation (SD). Chi-square test was used to compare categorical variables.

To determine differences in GEH between men and women, we constructed two linear regression models with sex as a predictor and normally distributed GEH variables (one-by-one) as an outcome. <u>Model 1</u> was adjusted for age and combinations of race and study center. To determine whether sex differences in GEH can be explained by sex differences in clinical and traditional ECG characteristics, <u>Model 2</u> was additionally adjusted for prevalent cardiovascular (CV) disease (HF, CHD, stroke), known CV risk factors (diabetes, hypertension, postmenopausal state in women, current smoking and alcohol intake, leisure physical activity level, levels of total cholesterol, high density lipoprotein (HDL), and triglycerides, BMI), use of antihypertensive and antiarrhythmic medications, serum concentrations of sodium, potassium, calcium, magnesium, phosphorus, and uric acid, total protein and albumin, blood urea nitrogen, CKD stage classified by eGFRCKD-EPI, education level, and traditional ECG characteristics [mean heart rate, QRS duration, Bazett-corrected QT interval, Cornell voltage, and sex-specific ECG – LVH].

#### Analysis of circular variables

Spatial QRS-T angle, SVG azimuth, and SVG elevation are circular variables. By convention, QRS-T and SVG elevation angles can be only positive, ranging from 0 to 180 degrees. Distributions of QRS-T angle and SVG elevation angle were normal or nearly normal. Thus, QRS-T and SVG elevation angles were included in all conventional statistical analyses without transformation. The SVG azimuth angle is expressed as an axial variable, ranging from - 180° to +180°. As recommended for the circular statistics^25^, we transformed SVG azimuth by doubling its value and then adding 360°. Then we analyzed the SVG azimuth using a conventional statistical approach, and for interpretation, we transformed it back.

#### Survival analyses

For an adequate comparison of separate GEH measurements, we assessed the hazard of SCD per 1 SD of continuous GEH variables, one-by-one. Similar models were constructed for traditional global ECG variables, one-by-one: heart rate, QRS duration, QTc, and Cornell voltage. Cox proportional hazards and Fine-Gray competing risks models were constructed. The proportional-hazards assumption was tested based on Schoenfeld residuals, using *stcox PH-assumptions* suite of tests implemented in STATA (StataCorp LP, College Station, TX). To adjust for possible confounders, we constructed <u>three models</u>, performed a statistical test for interaction with sex in each model, and constructed sex-stratified Cox models for men and women. The proportional-hazards assumption was confirmed for all predictors of interest in most models. Exceptions were reported. Relative hazard ratio (RHR) with a 95% confidence interval (CI) of SCD risk for women relative to men was reported, assuming HR for men is a reference (equal to 1).

<u>Model 1</u> was adjusted for: age and combinations of race and study center, prevalent HF, CHD, stroke, diabetes, hypertension, postmenopausal state, education level, current smoking, alcohol intake, leisure physical activity level, BMI category, use of antihypertensive and antiarrhythmic medications, levels of total cholesterol, HDL, and triglycerides, serum concentrations of sodium, potassium, calcium, magnesium, phosphorus, and uric acid, total protein and albumin, blood urea nitrogen, CKD stage classified by eGFRCKD-EPI, and sex-specific ECG-LVH. To avoid collinearity, models for Cornell voltage were not adjusted for ECG-LVH. Associations of continuous ECG variables with SCD were also evaluated using adjusted (model 1) Cox regression models incorporating cubic splines with 4 knots. The positions of the 4 knots in the cubic spline models are reported in Supplemental Table 1.

To determine whether global ECG measures associated with SCD independently from the dynamic substrate of structural heart disease, t<u>ime-updated model 2</u> included time-updated ECG predictors (one-by-one), all baseline covariates included in model 1, and time-updated incident nonfatal CVD (AF, HF, CHD, and stroke).

In addition, to determine whether GEH is associated with SCD independently from time-updated traditional ECG measures, <u>time-updated ECG-adjusted model 3</u> included time-updated GEH metrics (one-by-one), all four time-updated traditional ECG measurements (heart rate, QTc, QRS, and Cornell voltage), baseline clinical covariates, and time-updated incident nonfatal CVD included in model 2.

To study competing risks of SCD and nonSCD, we constructed Fine and Gray’s competing risks models for SCD and nonSCD outcomes, using the same covariates as described above for Cox models 1 and 2. Relative sub-hazard ratio (RSHR) with 95% CI of SCD risk for women relative to men was reported, assuming SHR for men is a reference.

Statistical analyses were performed using STATA MP 15.1 (StataCorp LP, College Station, TX). Considering the many multivariate analyses performed, statistical significance at the 0.05 level should be interpreted cautiously.

## Results

### Study population

Women comprised more than half of the study population (Table 1). Greater than half of the women were postmenopausal. At baseline, women had a lower prevalence of CVD as compared to men. Men had less favorable lipid profiles, were more likely current smokers and alcohol users, and were less physically active. However, there was a similar prevalence of diabetes and hypertension in men and women. There were significant differences in electrolytes and kidney function between men and women. Women had a faster heart rate, longer QTc, and a narrower QRS.

**Table 1.** Comparison of baseline clinical characteristics in men and women

| Characteristics | Men (n=6,601) | Women(n=8,124) | P-value |
| --- | --- | --- | --- |
| Age(SD), y | 54.6(5.8) | 53.8(5.7) | <0.0001 |
| White, n(%) | 5,229(78.1) | 5,886(71.4) | <0.0001 |
| Postmenopause, n(% of women) | n/a | 4,834(59.5) | n/a |
| Heart Failure, n(%) | 204(3.1) | 475(5.9) | <0.0001 |
| Coronary heart disease, n(%) | 528(8.0) | 169(2.1) | <0.0001 |
| Stroke, n(%) | 142(2.2) | 107(1.3) | <0.0001 |
| Body mass index(SD), kg/m<sup>2</sup> | 27.5(4.2) | 27.8(6.1) | 0.0002 |
| Diabetes, n(%) | 784(12.0) | 948(11.7) | 0.697 |
| Hypertension, n(%) | 2,227(33.7) | 2,811(34.6) | 0.272 |
| Antihypertensive drugs, n(%) | 1,782(27.0) | 2,664(32.8) | <0.0001 |
| Current tobacco smoker, n(%) | 1,809(27.4) | 2,020(24.9) | <0.0001 |
| Current alcohol drinker, n(%) | 4,282(64.9) | 4,010(49.4) | <0.0001 |
| Leisure physical activity score(SD) | 2.34(0.56) | 2.38(0.59) | 0.0001 |
| Education less than high school, n(%) | 1,543(23.4) | 1,863(22.9) | 0.526 |
| Total cholesterol(SD), mmol/L | 5.46(1.03) | 5.64(1.12) | <0.0001 |
| HDL cholesterol(SD), mg/dL | 44.3(13.8) | 57.6(17.3) | <0.0001 |
| Triglycerides(SD), mmol/L | 1.60(1.13) | 1.39(0.92) | <0.0001 |
| Sodium(SD), mmol/L | 140.8(2.4) | 141.0(2.5) | <0.0001 |
| Potassium(SD), mmol/L | 4.49(0.46) | 4.37(0.49) | <0.0001 |
| Calcium(SD), mg/dL | 9.76(0.42) | 9.81(0.44) | <0.0001 |
| Magnesium(SD), mEq/L | 1.64(0.16) | 1.63(0.16) | <0.0001 |
| Phosphorus(SD), mg/dL | 3.26(0.46) | 3.57(0.48) | <0.0001 |
| Total protein(SD), mg/dL | 7.27(0.44) | 7.28(0.46) | 0.024 |
| Albumin(SD), mg/dL | 3.92(0.26) | 3.83(0.27) | <0.0001 |
| Blood urea nitrogen(SD), mg/dL | 16.1(4.3) | 14.5(4.3) | <0.0001 |
| Chronic kidney disease stage≥2, n(%) | 2,310(35.0) | 2,247(27.7) | <0.0001 |
| Uric acid(SD), mg/dL | 6.73(1.42) | 5.48(1.43) | <0.0001 |
| Use of antiarrhythmic drugs, n(%) | 1,006(15.2) | 1,043(12.8) | <0.0001 |
| Heart rate(SD), bpm | 64.6(10.2) | 67.5(10.0) | <0.0001 |
| QRS duration(SD), ms | 96.9(12.5) | 88.4(10.7) | <0.0001 |
| QTc(SD), ms | 411.6(17.0) | 420.0(20.0) | <0.0001 |
| Cornell voltage(SD), μV | 1,403(588) | 1,103(495) | <0.0001 |
| Sex-specific ECG-LVH, n(%) | 423(6.4) | 419(5.2) | 0.001 |

2 HDL=High-density lipoprotein; SD=standard deviation

### Differences in GEH between men and women

In both unadjusted and adjusted analyses, QRS-T angle and SAI QRST were significantly larger in men as compared to women. (Table 2 and Supplemental Figure 1). In contrast, sex differences in SVG magnitude were explained by covariates. SVG vector pointed more upward and forward in men, and the difference in SVG direction not only remained significant after full adjustment but increased up to 15-17 degrees.

**Table 2.** Difference in GEH variables in women as compared to men

| GEH characteristic | Model 1 |  | Model 2 |  |
| --- | --- | --- | --- | --- |
|  | Difference (95%CI) | P-value | Difference (95%CI) | P-value |
| Peak QRS-T angle, ° | -12.1(-13.1 to -11.0) | <0.0001 | -8.2(-10.7 to -5.7) | <0.0001 |
| Area QRS-T angle, ° | -15.5(-16.4 to -14.6) | <0.0001 | -9.5(-11.6 to -7.5) | <0.0001 |
| Peak SVG elevation, ° | -5.95(-6.43 to -5.46) | <0.0001 | -2.33(-3.43 to -1.22) | <0.0001 |
| Area SVG elevation, ° | -5.01(-5.56 to -4.46) | <0.0001 | -3.42(-4.74 to -2.10) | <0.0001 |
| Peak SVG azimuth, ° | +11.27(+9.65 to +12.88) | <0.0001 | +13.58(+9.72 to +17.44) | <0.0001 |
| Area SVG azimuth, ° | +8.94(+7.32 to +10.57) | <0.0001 | +11.95(+8.16 to +15.74) | <0.0001 |
| SAI QRST, mV*ms | -33.6(-35.1 to -32.0) | <0.0001 | -12.1(-15.4 to -8.8) | <0.0001 |
| Peak SVG magnitude, μV | -51.6(-65.3 to -37.8) | <0.0001 | +47.6(+12.9 to + 82.3) | 0.007 |
| SVG magnitude, μV | -92.7(-107.9 to -77.5) | <0.0001 | -14.8(-52.2 to -22.7) | 0.439 |

**Model 1** was adjusted for age and combination of race and study center. **Model 2** was in
addition adjusted for prevalent heart failure, coronary heart disease, stroke, diabetes,
hypertension, body mass index, postmenopause state, education level, current smoking, current
alcohol intake, leisure physical activity level, use of antihypertensive and antiarrhythmic
medications, levels of total cholesterol, high density lipoprotein, and triglycerides, serum
concentrations of sodium, potassium, calcium, magnesium, phosphorus, and uric acid, level total
protein and albumin, blood urea nitrogen, chronic kidney disease stage classified by eGFR<sub>CKD-</sub>
EPI, mean heart rate, QRS duration, corrected QT interval, Cornell voltage, and sex-specific ECG
– left ventricular hypertrophy.

### EP substrate of sudden cardiac death in men and women in Cox regression analysis

Over a median follow-up of 24.4 years, there were 530 SCDs (incidence 1.72; 95% CI 1.58-1.88 per 1000 person-years), 2,178 nonSCDs (incidence 7.09; 95% CI 6.80-7.39 per 1000 person-years), and 2,535 noncardiac deaths (incidence 8.25; 95%CI 7.93-8.58 per 1000 person-years). Incidence of SCD was higher in men (2.56; 95%CI 2.30-2.84 per 1000 person-years) than in women (1.10; 95%CI 0.95-1.26 per 1000 person-years). Incidence of nonSCD was also higher in men (8.51; 95%CI 8.03-9.03 per 1000 person-years) than in women (6.01; 95%CI 5.66-6.38 per 1000 person-years). Similarly, noncardiac death was also more frequent in men (incidence 10.51; 95% CI 9.97-11.08 per 1000 person-years) than in women (incidence 6.54; 95% CI 6.17-6.93 per 1000 person-years).

In Cox model 1, QRS-T angle, SVG direction, heart rate, and Cornell voltage were associated with SCD (Figure 1 and Supplemental Table 2). Further adjustment for time-updated CHD, HF, AF, and stroke strengthened the association of nearly all ECG metrics with SCD. In Cox model 2, all studied ECG metrics, except SVG magnitude, were associated with SCD.

**Figure 1.**
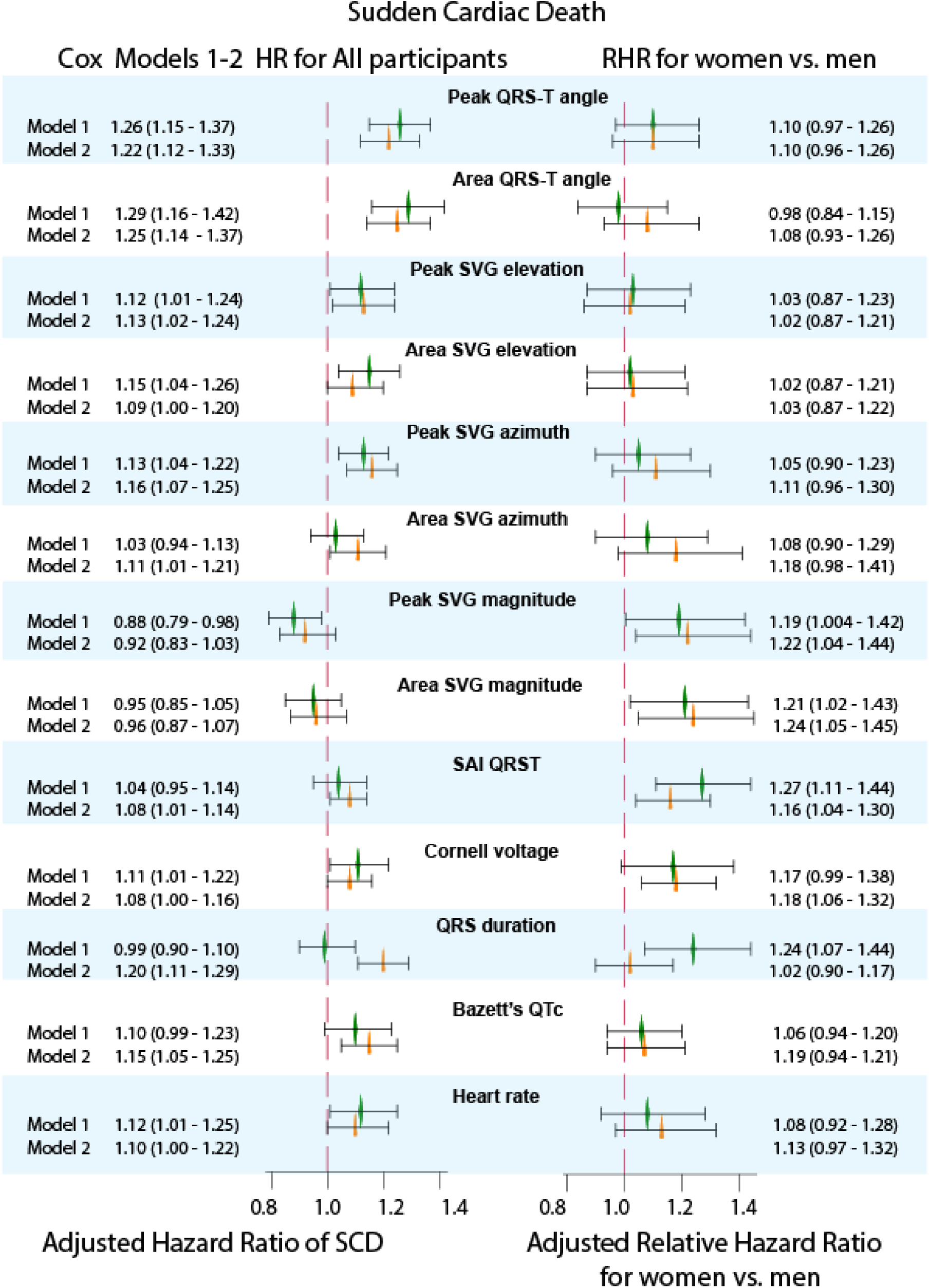
Adjusted Cox proportional hazard ratio (HR) and 95% confidence interval (CI) of SCD for GEH and traditional global ECG metrics in model 1 (green diamond) and model 2 (orange triangle). Black lines correspond to 95% CI bounds. Left forest plot shows HR with 95%CI for all participants. Right forest plot shows relative HR (RHR) with 95%CI for women as compared to men, with HR for men equal 1.0.

We observed a statistically significant interaction of sex with SAI QRST, SVG magnitude, and QRS duration (Figure 1 and Supplemental Table 2). In model 1, there was a 19-27% higher risk of SCD in women compared to men, per one SD of SVG magnitude, SAI QRST, and QRS duration. Adjustment for incident nonfatal CVD in model 2 further strengthened the interaction of sex with SVG magnitude and revealed significant interaction with Cornell voltage. However, model 2 attenuated the interaction with SAI QRST and wiped out the interaction with QRS duration.

Sex-stratified Cox models confirmed a significant association of traditional and novel global ECG metrics with SCD (Figure 2 and Supplemental Table 2B). Larger SVG magnitude pointed towards a higher risk of SCD in women. In contrast, a larger SVG magnitude trended towards a lower risk of SCD in men. The strength of the association of SVG magnitude with SCD did not reach statistical significance, but opposite trends were seen (Supplemental Figure 2H). After full adjustment for nonfatal incident CVD, there was a 24% increase SCD risk in women versus 10% in men with one SD increase in Cornell voltage. Similarly, there was a 19% increase in SCD risk in women versus 9% in men with one SD of SAI QRST.

**Figure 2.**
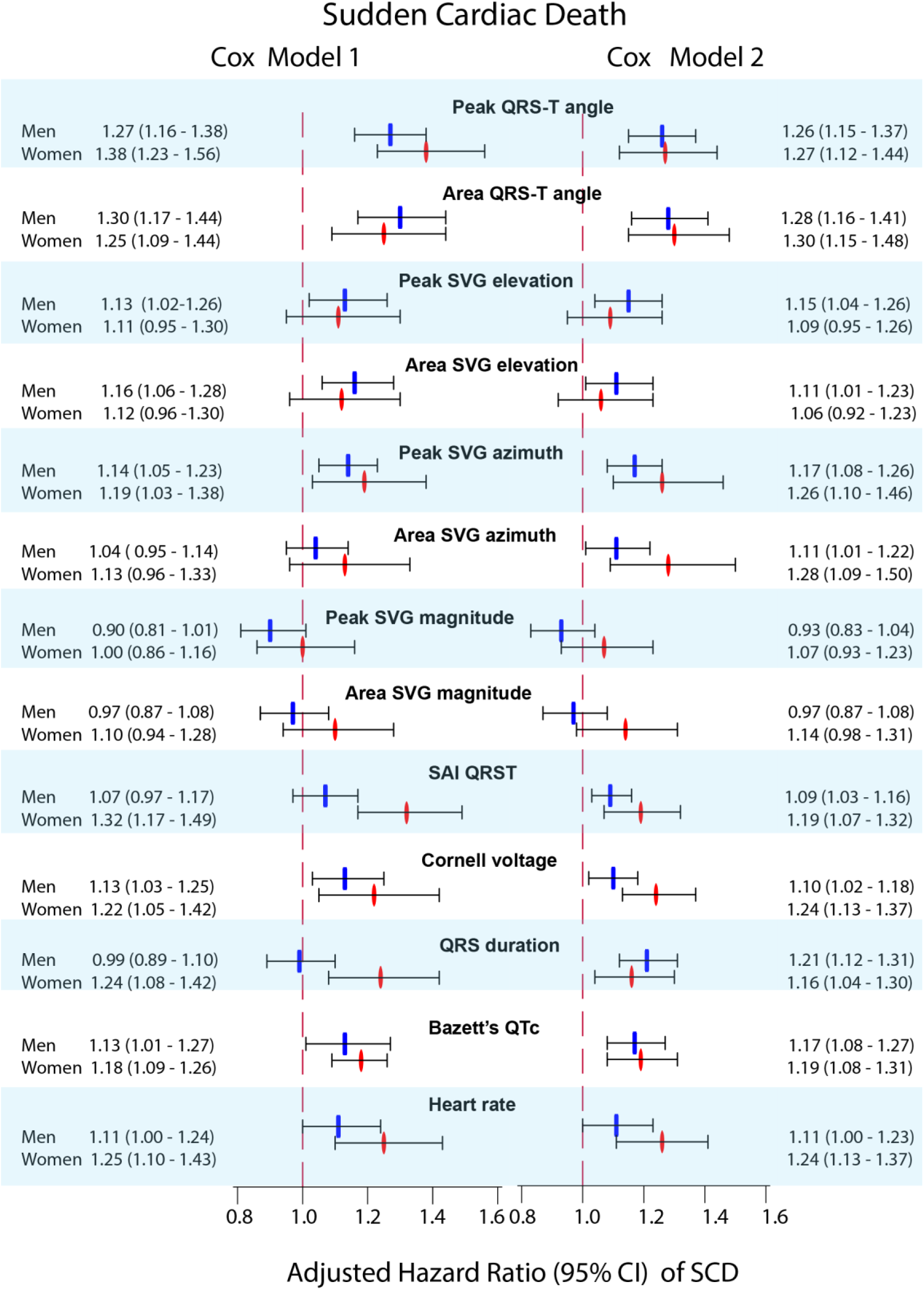
Sex-stratified adjusted (models 1 and 2) Cox proportional hazard ratio (HR) and 95% confidence interval (CI) of SCD for GEH and traditional global ECG metrics in men (blue rectangle) and women (red oval). Black lines correspond to 95% CI bounds.

Interaction of SVG magnitude and SAI QRST with sex remained significant after further adjustment for time-updated traditional ECG metrics (heart rate, QTc, QRS, and Cornell voltage) in model 3. In women, greater SVG magnitude was associated with a higher risk of SCD, whereas in men, bigger SVG magnitude and SAI QRST tended to be protective (Supplemental Table 2B).

### Competing risks of SCD and nonSCD

In a competing risk model 1, one SD increase in spatial QRS-T angle or SVG direction (azimuth and elevation) was associated with a 10-19% increase in odds of SCD occurrence (Supplemental Table 3A and Figure 3). Traditional ECG metrics were not associated with SCD in the competing risks analysis. Competing risk model 2 only slightly attenuated the association of QRS-T angle, SVG elevation, and SVG azimuth with SCD, and revealed a significant association of QRS duration with SCD.

**Figure 3.**
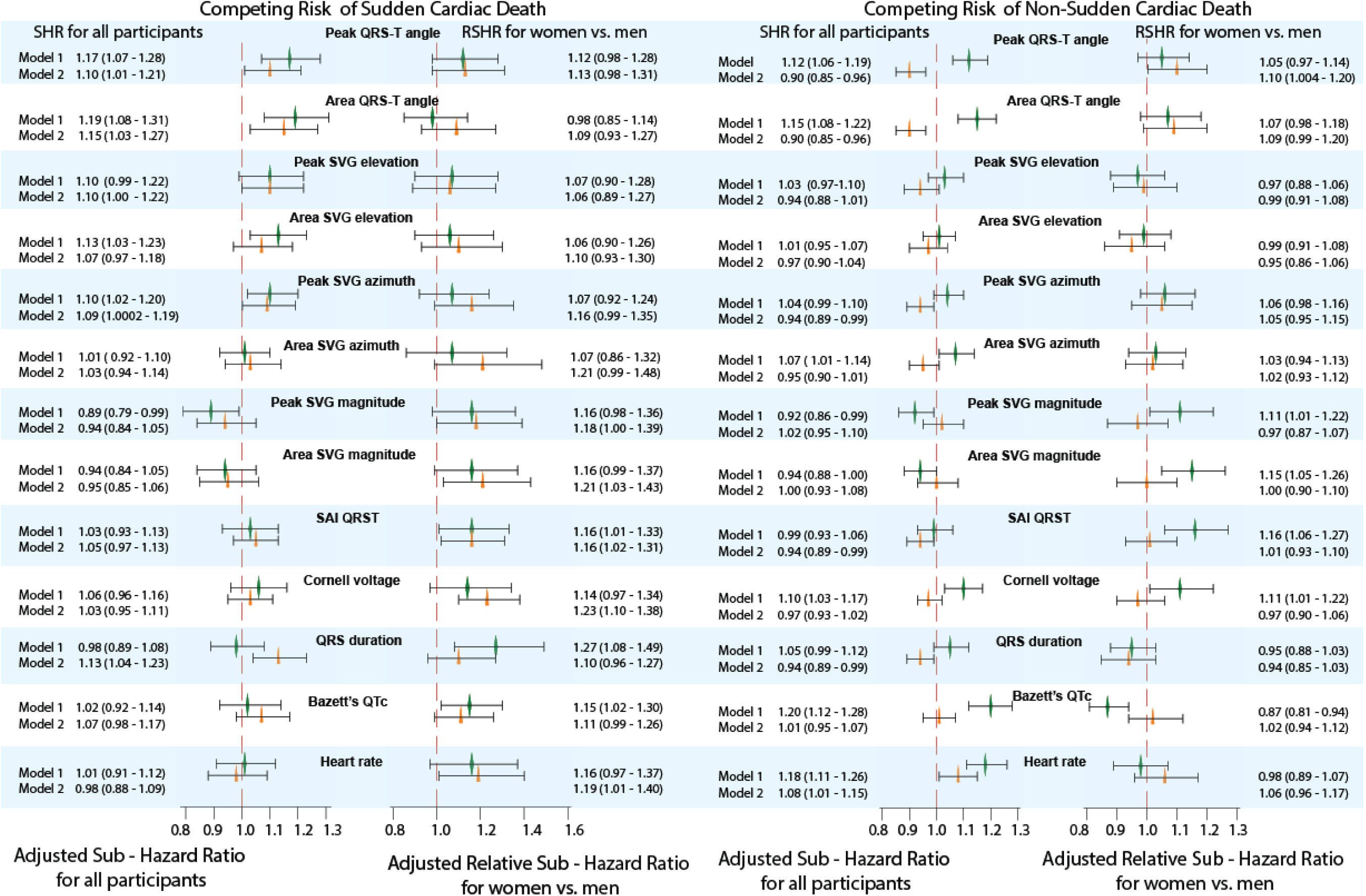
Adjusted competing risk sub-hazard ratio (SHR) and 95% confidence interval (CI) of SCD for GEH and traditional global ECG metrics in model 1 (green diamond) and model 2 (orange triangle). Black lines correspond to 95% CI bounds. Left forest plot shows SHR with 95%CI for all participants. Right forest plot shows relative SHR (RSHR) with 95%CI for women as compared to men, with SHR for men equal 1.0.

In competing risk model 1, QRS-T angle, SVG azimuth, Cornell voltage, QTc, and heart rate were associated with an increased incidence of nonSCD (Figure 3 and Supplemental Table 3A). Of note, greater SVG magnitude was associated with *decreased* incidence of nonSCD. Incident nonfatal CVD explained the association of QTc and Cornell voltage with nonSCD, whereas heart rate remained independently associated with nonSCD even after adjustment in model 2. Of note, after adjustment in model 2, SAI QRST, QRS duration, QRS-T angle, and SVG magnitude were associated with *decreased* incidence of nonSCD, mirroring observed *increased* incidence of SCD associated with these ECG metrics in competing risk model 2 for SCD.

### Relative competing risk of SCD and nonSCD in women as compared to men

In competing risk model 1, a statistically significant interaction of sex with competing risk of SCD was observed for QTc, QRS duration, and SAI QRST. Women experienced a greater increase in odds of SCD occurrence compared to men: by 27% per SD of QRS duration, 16% per SD of SAI QRST, and 15% per SD of QTc interval.

Adjustment for dynamic CVD substrate eliminated the interaction with QTc and QRS duration, suggesting that sex differences in SCD risk conveyed by QTc and QRS duration were explained by sex differences in structural heart disease substrate. Model 2, however, revealed significant interaction of sex with SVG magnitude and Cornell voltage, in addition to interaction with SAI QRST. After full adjustment for incident CVD, SVG magnitude, SAI QRST, and Cornell voltage were associated with 16-23% increase in odds of SCD occurrence in women as compared to men (Figure 3 and Supplemental Table 3A).

A few interactions were observed for competing risk of nonSCD in model 1, but not in model 2. This suggests that sex differences in the risk of nonSCD were explained by incident nonfatal CVD.

In sex-stratified analyses (Figure 4 and Supplemental Table 3B), in adjusted for baseline confounders model 1, QTc, QRS, and SAI QRST were associated with increased odds of SCD occurrence by 18-26% in women, but not in men. In men, but not in women, QTc prolongation and smaller peak SVG magnitude were associated with an increased incidence of nonSCD.

**Figure 4.**
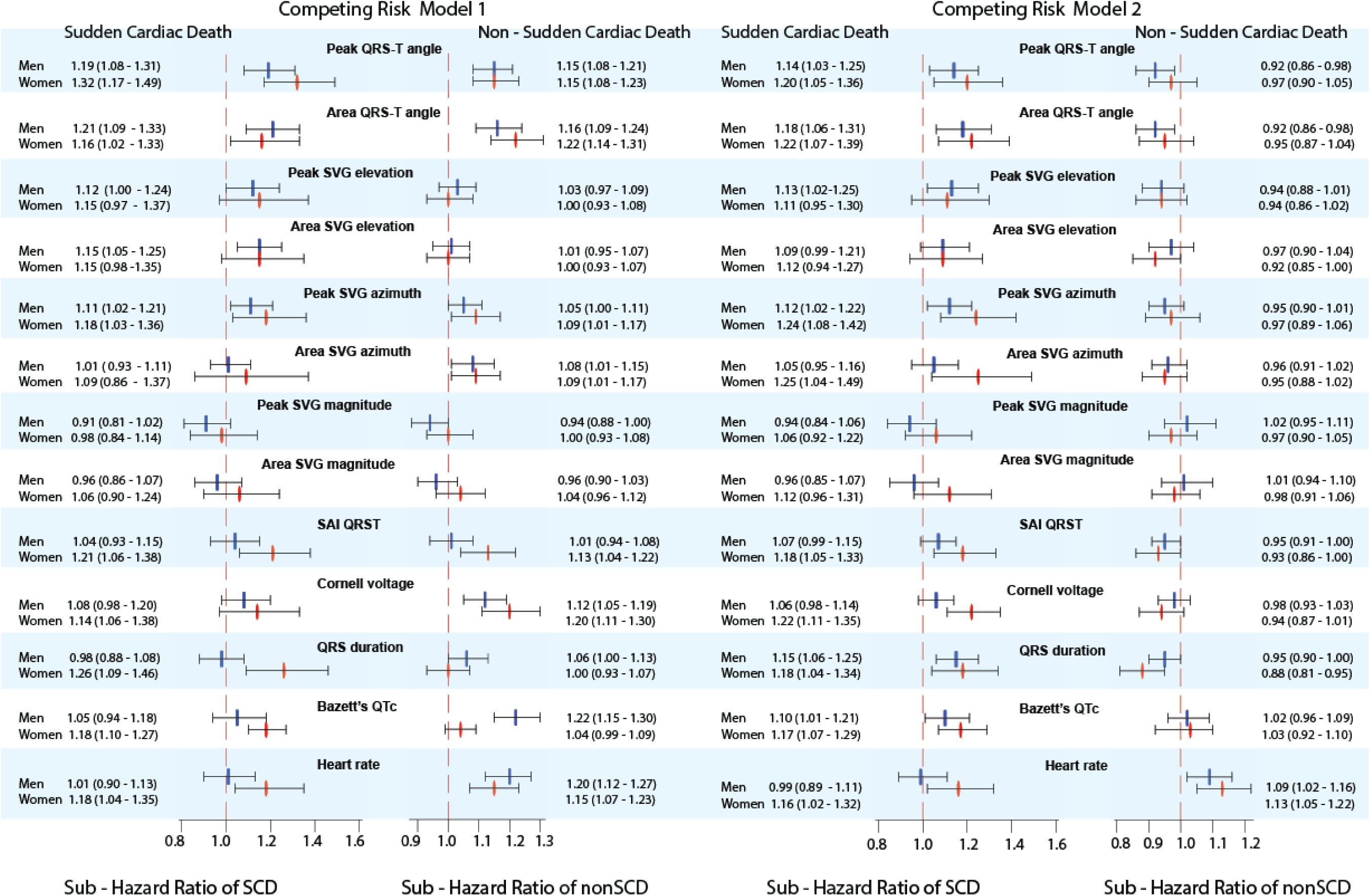
Sex-stratified adjusted (models 1 and 2) competing risk sub-hazard ratio (SHR) and 95% confidence interval (CI) of SCD and nonSCD for GEH and traditional global ECG metrics in men (blue rectangle) and women (red oval). Black lines correspond to 95% CI bounds.

When adjusted for dynamic CVD substrate in model 2, in women, larger SAI QRST, QRS duration, SVG magnitude, and Cornell voltage were associated with greater odds of SCD. As expected in mirroring competing risk model, smaller SAI QRST, QRS duration, SVG magnitude, and Cornell voltage were associated with an increased incidence of nonSCD.

Across all comparisons and models, peak-based and area-based GEH metrics displayed consistent results, reassuring robustness of analyses.

## Discussion

Our study of a large, community-based prospective cohort of over 14,000 participants with greater than 24 years median follow-up showed that sex is a significant modifier with respect to the association of EP substrate with SCD (Figure 5). In women, global EP substrate (QRS duration, Cornell voltage, SAI QRST, SVG magnitude, heart rate, and QTc) was associated with up to 27% greater risk of SCD than in men. Our findings have important clinical implications: development of sex-specific risk score of SCD is necessary, and the addition of global EP substrate metrics in the risk prediction model for women is warranted. Further studies of mechanisms behind global EP substrate in men and women are needed for the development of sex-specific prevention of SCD. Theoretically, there are two major groups of mechanisms behind the observed effect modification: differences in the cardiac EP substrate between men and women, and differences in structural heart disease substrate.

**Figure 5.**
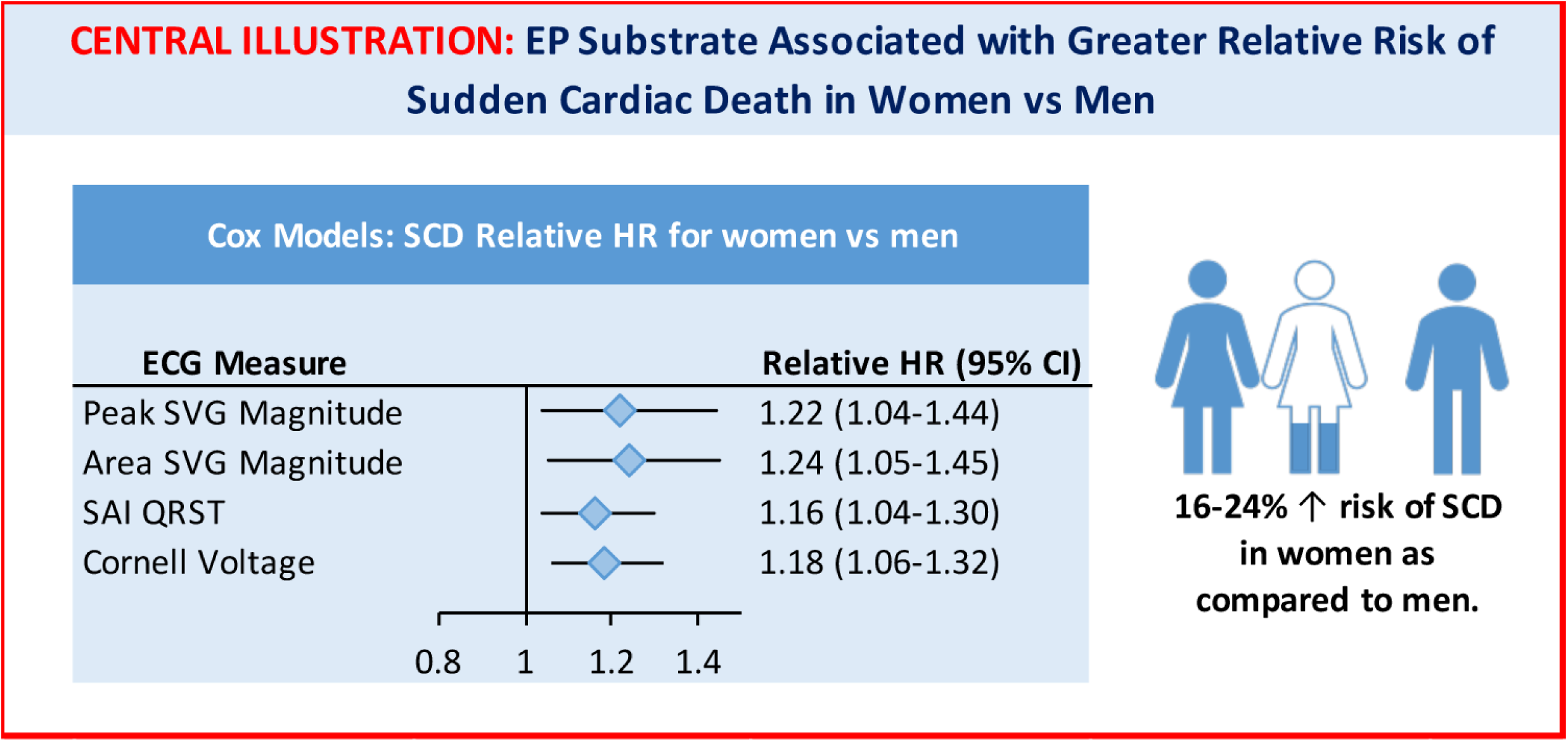
Summary of findings

### Why does EP substrate associated with greater risk of SCD in women? EP hypothesis

Our study showed that after rigorous adjustment for baseline demographic and clinical risk factors of SCD, including prevalent CVD and CV risk factors, postmenopausal state, serum concentrations of electrolytes and degree of CKD, several traditional ECG metrics (QRS duration, heart rate, and QTc), Cornell voltage, and voltage-based GEH metrics (SAI QRST and SVG magnitude) were associated with greater SCD risk in women than in men. In men, EP substrate was explained by an underlying CVD, whereas in women, EP substrate conveyed an additional risk of SCD, beyond the risk carried by the prevalent CVD and CV risk factors.

The most remarkable difference in the risk of SCD between men and women was conveyed by amplitude-based ECG metrics: Cornell voltage, SAI QRST, and SVG magnitude. Importantly, the interaction of sex with amplitude-based ECG metrics was independent not only from baseline CVD and its risk factors but also from incident CVD, and it was consistently observed in both Cox regression analysis and competing risk models. One SD increase in Cornell voltage was associated with more than 20% higher risk of SCD in women as compared to men. Our finding is consistent with a recent autopsy SCD study in the Finnish population, which observed ECG-LVH more commonly in female than male SCD victims.^26^

We observed that one SD increase in the magnitude of SVG (expressed either as SVG vector magnitude, or SVG’s scalar, SAI QRST) was associated with approximately 20% higher risk of SCD in women as compared to men. A recent Finnish study demonstrated results consistent with our findings of sex differences in SAI QRST and its association with fatal CVD,^27^ although it did not specifically include SCD. The magnitude of SVG and SAI QRST are global measures of the dispersion of total recovery time in the heart, encompassing dispersion of activation and refractoriness.^28^ Women have greater asymmetry in potassium channel expression between left and right ventricles.^29^ In a recent genome-wide association study, SAI QRST and SVG magnitude were associated with genetic polymorphisms tagging *HAND1* and *TBX3* genes, involving mechanisms of left to right asymmetry in the heart.^30^ We speculate that SVG magnitude and SAI QRST reflect differences in cardiac electrophysiology between men and women, which are responsible for the stronger association of SAI QRST and SVG magnitude with SCD in women than in men.

We demonstrated that QRS duration is associated with more than 20% higher SCD risk in women than in men, as demonstrated by both Cox regression and competing risks analyses. Sex differences in SCD risk conveyed by QRS duration were largely explained by sex differences in dynamic structural heart disease substrate. Existing literature on the association between QRS duration and SCD is inconsistent, likely owing at least in part to the study populations having very few women (1-16%) and the majority of analyses lacking stratification by sex.^31–33^ Similar mechanisms may be responsible for why women derive greater benefit from cardiac resynchronization therapy which remains incompletely understood.^9^

### Comparison of Cox proportional hazards and Fine-Gray competing risk regression results

SCD and nonSCD events are naturally competing, tightly intertwined events and cannot be studied in isolation. CVD continuum encompasses progression from CVD risk factors to subclinical and then to clinically manifested CVD, and, ultimately to either SCD or nonSCD. To develop a greater understanding of relationships between EP substrate and SCD and effect modification by sex, we fitted both Cox regression and competing risk models, and appropriately interpreted the regression coefficients from the subdistribution hazard model.^34^ It was previously shown that when the probability of an event is less than 0.2, the logistic link function and the complementary log-log link function are very similar,^34^ and a subdistribution hazard model can be interpreted as odds ratios for the cumulative incidence function. In this study, the probability of SCD, but not a probability of nonSCD met these criteria. In this study, voltage-based ECG metrics (SVG magnitude, SAI QRST, Cornell voltage) and QRS duration demonstrated greater risk of SCD for women as compared to men in both Cox and Fine-Gray models. However, QTc and heart rate were stronger associated with SCD in women than in men in competing risk models only, but not in Cox models. Statistically significant interactions with sex revealed in Fine-Gray models highlight the importance of competing risk analysis for understanding complex relationships of EP substrate with SCD and nonSCD in men and women.

Our study showed that in women, QTc is associated with greater odds of SCD, whereas in men, QTc is associated with greater incidence of nonSCD. While QT prolongation is a known risk marker of torsades de pointes (TdP) in congenital long QT syndrome,^35^ in other populations, the association of QT interval with SCD was controversial.^36^ One possible reason for controversy around the association of QTc with SCD can be explained by differences in the proportion and clinical characteristics of women enrolled in previous studies. No prior studies tested statistical interaction of sex with QTc after extensive adjustment for confounders. Consistently with our findings, the Rotterdam study showed an association of QT prolongation with SCD only in the absence of cardiac dysfunction, whereas, in patients with systolic HF, risk of SCD was independent of QT prolongation.^37^ Similarly, OregonSUDS study reported the stronger association of QTc prolongation with SCD in diabetes-free individuals as compared to those with diabetes.^38^ Women have a longer QT interval due to reduced expression of potassium channels, resulting in decreased rapid and slow delayed rectifier K^+^ currents, inward rectifier current, and transient outward current.^39, 40^ Estrogens inhibit the rapid delayed rectifier current, increase the L-type calcium current, the sodium-calcium exchange current, and calcium release mediated by the ryanodine receptor, which can predispose to triggered activity.^41^ Two-thirds of the drug-induced TdP cases occur in women.^42^ Thus, in women, QTc carries additional risk of SCD due to sex-specific EP mechanisms, independent of common for men and women CVD substrate.

In this study, resting heart rate was associated with greater odds of SCD in women but not in men. Association of a resting heart rate with SCD in women was independent of incident CVD, supporting previous OregonSUDS findings.^43^ Women have faster resting heart rate^9^ mostly because of smaller LV mass and volume, resulting in lesser exercise capacity in women than in men.^44^ Exercise capacity is associated with cardiac arrhythmias.^45^ Our results suggest that lesser exercise capacity in women, manifesting by faster resting heart rate, translates into the stronger association of heart rate with SCD in women, which is independent of the CVD development.

### Sex differences in structural heart disease substrate

In this study, non-fatal incident CVD explained the stronger association of QTc and QRS duration with SCD in women, as compared to men. On another hand, sex did not modify the association of studied ECG features with nonSCD. This finding is in accord with known differences in structural heart disease between men and women. In spite of less frequent obstructive CHD, women with angina or MI have greater cardiac mortality than men.^46, 47^ Women have different coronary microvasculature and greater arteriolar wall thickness than men.^48^ On the other hand, men are more likely to develop cardiac amyloidosis (manifesting by small ECG voltage), and subsequently HF.^49^ Thus, in women, QTc and QRS duration reflect an underlying structural heart disease with greater than in men risk of proarrhythmia, whereas, in men, QTc and QRS duration reflect an underlying structural heart disease leading to pump failure and eventually, more likely to nonSCD.

### Differences in GEH between men and women

Consistent with previous studies in healthy young individuals^50^ and young athletes^12^, we observed wider QRS-T angle, larger SAI QRST, and SVG vector pointing more upward and forward in middle-aged men than in middle-aged women. Of note, our study revealed that differences in SVG magnitude between men and women are explained by differences in body size, and other clinical characteristics, both cardiac and non-cardiac, suggesting that fundamentally, there is no difference in the magnitude of gradient between the longest and the shortest action potential duration between male and female heart.

### Clinical implications of greater risk of SCD associated with global EP substrate in women

We observed the significantly stronger association of several ECG metrics of underlying EP substrate (QRS duration, Cornell voltage, SAI QRST, SVG magnitude, heart rate, and QTc) with SCD in women than in men. Therefore, the addition of these ECG metrics in the risk prediction model for women is warranted for the development of a sex-specific risk score of SCD in women. Our results indicate that significant improvement in SCD risk prediction for women can be made. Improvement of SCD risk stratification is especially important for women considering primary prevention ICD.^4^ Further studies of sex-specific EP substrate in men and women are needed for the development of future therapies.

### Strengths and Limitations

This is a large community-based prospective cohort study with long-term follow-up, well-adjudicated SCD, and approximately equal representation of men and women, providing unique opportunity to study sex exposure as an effect modifier. The well-characterized population of the ARIC study allowed us to perform comprehensive adjustment for confounders, including post-menopausal state, electrolytes, and kidney function, accounting for important non-cardiac differences between men and women. However, limitations of the study have to be taken into account. The study population was predominantly white; only 26% of the study participants were black. Validation of the study finding in a multiracial population is needed. Due to the lack of information on baseline LVEF for most of the study participants, we did not adjust our analyses for baseline LVEF. Nevertheless, we adjusted our analyses for incident HF and conducted competing risk analyses, sufficiently accounting for competing risk of a pump failure death.

## Acknowledgments

The authors thank the staff and participants of the ARIC study for their important contributions. We would like to acknowledge the SCD mortality classification committee members: Nona Sotoodehnia (lead), Selcuk Adabag, Sunil Agarwal, Lin Chen, Rajat Deo, Leonard Ilkhanoff, Liviu Klein, Saman Nazarian, Ashleigh Owen, Kris Patton, and Larisa Tereshchenko.

## SUPPLEMENTAL MATERIALS

### Supplemental Tables

**Supplemental Table 1.** The positions of knots in the cubic spline models

| ECG metric | Men (n=6,601) |  |  |  | Women(n=8,124) |  |  |  |
| --- | --- | --- | --- | --- | --- | --- | --- | --- |
|  | Knot #1 | Knot#2 | Knot#3 | Knot#4 | Knot #1 | Knot#2 | Knot#3 | Knot#4 |
| Peak QRS-T angle, ° | 12.5 | 33.4 | 53.2 | 138.0 | 8.9 | 25.6 | 40.7 | 110.2 |
| Area QRS-T angle, ° | 28.0 | 57.2 | 77.8 | 122.3 | 18.1 | 42.3 | 60.5 | 102.3 |
| Peak SVG elevation, ° | 41.7 | 61.4 | 71.5 | 92.3 | 38.3 | 55.8 | 65.7 | 82.4 |
| Area SVG elevation, ° | 44.1 | 62.3 | 74.5 | 104.9 | 42.5 | 58.6 | 69.7 | 93.2 |
| Peak SVG azimuth, ° | -55.7 | -11.4 | 1.8 | 69.8 | -19.6 | -1.6 | 9.6 | 42.3 |
| Area SVG azimuth, ° | -18.8 | 12.6 | 30.6 | 65.1 | -6.2 | 18.4 | 33.9 | 61.8 |
| SAI QRST, mV*ms | 92.2 | 135.4 | 171.0 | 257.5 | 79.2 | 110.8 | 135.5 | 195.2 |
| Peak SVG magnitude, µV | 919 | 1432 | 1764 | 2404 | 957 | 1407 | 1723 | 2332 |
| SVG magnitude, µV | 988 | 1517 | 1898 | 2676 | 1012 | 1484 | 1820 | 2473 |

**Supplemental Table 2A:**
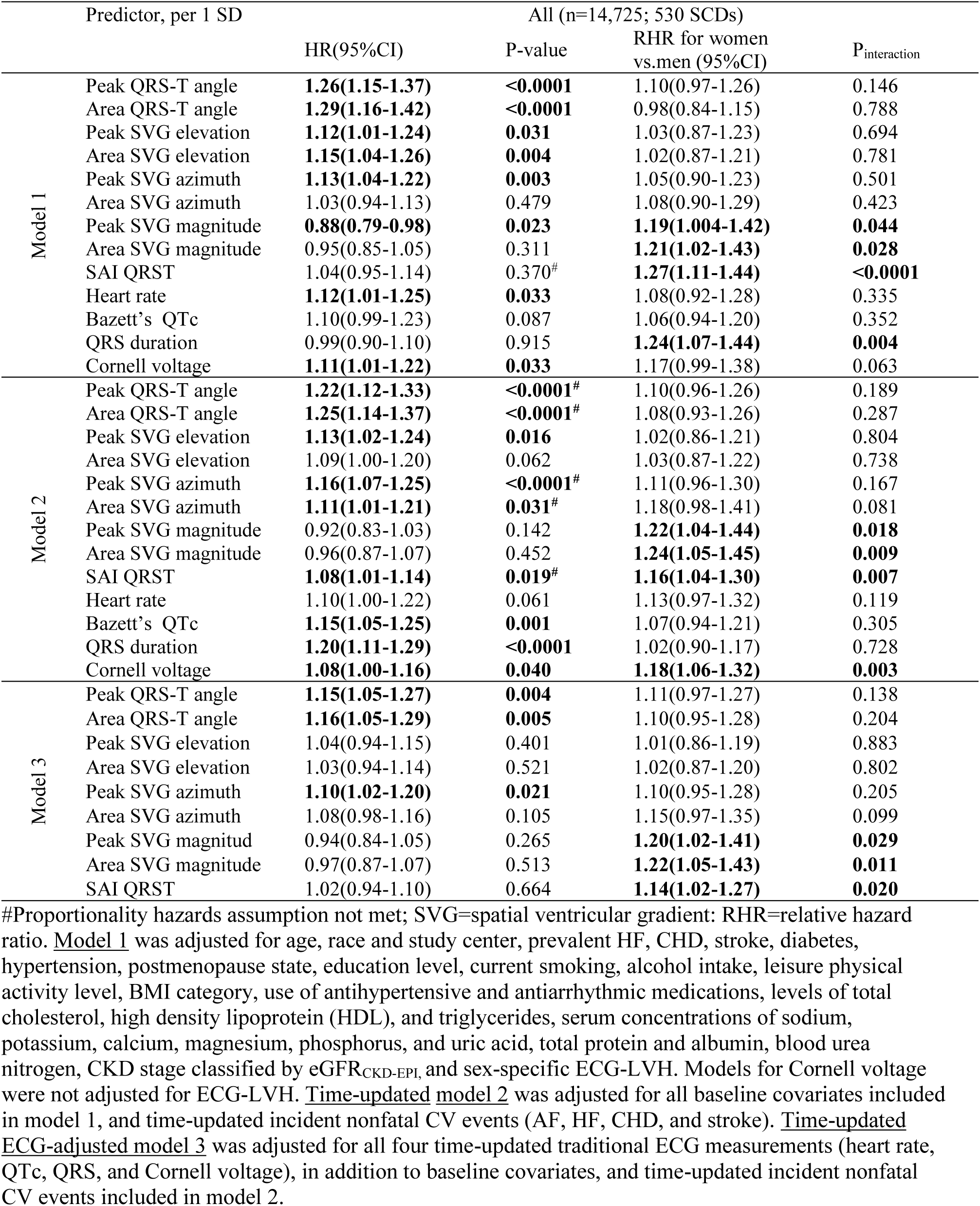
Sex interaction in association of GEH with SCD in Cox models

| Predictor, per 1 SD |  | All (n=14,725; 530 SCDs) |  |  |  |
| --- | --- | --- | --- | --- | --- |
|  |  | HR(95%CI) | P-value | RHR for women vs.men (95%CI) | P<sub>interaction</sub> |
| Model 1 | Peak QRS-T angle | **1.26(1.15-1.37)** | **<0.0001** | 1.10(0.97-1.26) | 0.146 |
|  | Area QRS-T angle | **1.29(1.16-1.42)** | **<0.0001** | 0.98(0.84-1.15) | 0.788 |
|  | Peak SVG elevation | **1.12(1.01-1.24)** | **0.031** | 1.03(0.87-1.23) | 0.694 |
|  | Area SVG elevation | **1.15(1.04-1.26)** | **0.004** | 1.02(0.87-1.21) | 0.781 |
|  | Peak SVG azimuth | **1.13(1.04-1.22)** | **0.003** | 1.05(0.90-1.23) | 0.501 |
|  | Area SVG azimuth | 1.03(0.94-1.13) | 0.479 | 1.08(0.90-1.29) | 0.423 |
|  | Peak SVG magnitude | **0.88(0.79-0.98)** | **0.023** | **1.19(1.004-1.42)** | **0.044** |
|  | Area SVG magnitude | 0.95(0.85-1.05) | 0.311 | **1.21(1.02-1.43)** | **0.028** |
|  | SAI QRST | 1.04(0.95-1.14) | 0.370<sup>#</sup> | **1.27(1.11-1.44)** | **<0.0001** |
|  | Heart rate | **1.12(1.01-1.25)** | **0.033** | 1.08(0.92-1.28) | 0.335 |
|  | Bazett's QTc | 1.10(0.99-1.23) | 0.087 | 1.06(0.94-1.20) | 0.352 |
|  | QRS duration | 0.99(0.90-1.10) | 0.915 | **1.24(1.07-1.44)** | **0.004** |
|  | Cornell voltage | **1.11(1.01-1.22)** | **0.033** | 1.17(0.99-1.38) | 0.063 |
| Model 2 | Peak QRS-T angle | **1.22(1.12-1.33)** | **<0.0001<sup>#</sup>** | 1.10(0.96-1.26) | 0.189 |
|  | Area QRS-T angle | **1.25(1.14-1.37)** | **<0.0001<sup>#</sup>** | 1.08(0.93-1.26) | 0.287 |
|  | Peak SVG elevation | **1.13(1.02-1.24)** | **0.016** | 1.02(0.86-1.21) | 0.804 |
|  | Area SVG elevation | 1.09(1.00-1.20) | 0.062 | 1.03(0.87-1.22) | 0.738 |
|  | Peak SVG azimuth | **1.16(1.07-1.25)** | **<0.0001<sup>#</sup>** | 1.11(0.96-1.30) | 0.167 |
|  | Area SVG azimuth | **1.11(1.01-1.21)** | **0.031<sup>#</sup>** | 1.18(0.98-1.41) | 0.081 |
|  | Peak SVG magnitude | 0.92(0.83-1.03) | 0.142 | **1.22(1.04-1.44)** | **0.018** |
|  | Area SVG magnitude | 0.96(0.87-1.07) | 0.452 | **1.24(1.05-1.45)** | **0.009** |
|  | SAI QRST | **1.08(1.01-1.14)** | **0.019<sup>#</sup>** | **1.16(1.04-1.30)** | **0.007** |
|  | Heart rate | 1.10(1.00-1.22) | 0.061 | 1.13(0.97-1.32) | 0.119 |
|  | Bazett's QTc | **1.15(1.05-1.25)** | **0.001** | 1.07(0.94-1.21) | 0.305 |
|  | QRS duration | **1.20(1.11-1.29)** | **<0.0001** | 1.02(0.90-1.17) | 0.728 |
|  | Cornell voltage | **1.08(1.00-1.16)** | **0.040** | **1.18(1.06-1.32)** | **0.003** |
| Model 3 | Peak QRS-T angle | **1.15(1.05-1.27)** | **0.004** | 1.11(0.97-1.27) | 0.138 |
|  | Area QRS-T angle | **1.16(1.05-1.29)** | **0.005** | 1.10(0.95-1.28) | 0.204 |
|  | Peak SVG elevation | 1.04(0.94-1.15) | 0.401 | 1.01(0.86-1.19) | 0.883 |
|  | Area SVG elevation | 1.03(0.94-1.14) | 0.521 | 1.02(0.87-1.20) | 0.802 |
|  | Peak SVG azimuth | **1.10(1.02-1.20)** | **0.021** | 1.10(0.95-1.28) | 0.205 |
|  | Area SVG azimuth | 1.08(0.98-1.16) | 0.105 | 1.15(0.97-1.35) | 0.099 |
|  | Peak SVG magnitud | 0.94(0.84-1.05) | 0.265 | **1.20(1.02-1.41)** | **0.029** |
|  | Area SVG magnitude | 0.97(0.87-1.07) | 0.513 | **1.22(1.05-1.43)** | **0.011** |
|  | SAI QRST | 1.02(0.94-1.10) | 0.664 | **1.14(1.02-1.27)** | **0.020** |

<sup>#</sup>Proportionality hazards assumption not met; SVG=spatial ventricular gradient; RHR=relative hazard ratio. Model 1 was adjusted for age, race and study center, prevalent HF, CHD, stroke, diabetes, hypertension, postmenopause state, education level, current smoking, alcohol intake, leisure physical activity level, BMI category, use of antihypertensive and antiarrhythmic medications, levels of total cholesterol, high density lipoprotein (HDL), and triglycerides, serum concentrations of sodium, potassium, calcium, magnesium, phosphorus, and uric acid, total protein and albumin, blood urea nitrogen, CKD stage classified by eGFR<sub>CKD-EPI</sub>, and sex-specific ECG-LVH. Models for Cornell voltage were not adjusted for ECG-LVH. Time-updated model 2 was adjusted for all baseline covariates included in model 1, and time-updated incident nonfatal CV events (AF, HF, CHD, and stroke). Time-updated ECG-adjusted model 3 was adjusted for all four time-updated traditional ECG measurements (heart rate, QTc, QRS, and Cornell voltage), in addition to baseline covariates, and time-updated incident nonfatal CV events included in model 2.

**Supplemental Table 2B:** Association of GEH with SCD in Cox models for men and women

| Predictor, per 1 SD |  | Men (n=6,601; 338 SCDs) |  | Women (n=8,124; 192 SCDs) |  |
| --- | --- | --- | --- | --- | --- |
|  |  | HR(95%CI) | P-value | HR(95%CI) | P-value |
| Model 1 | Peak QRS-T angle | **1.27(1.16-1.38)** | **<0.0001** | **1.38(1.23-1.56)** | **<0.0001** |
|  | Area QRS-T angle | **1.30(1.17-1.44)** | **<0.0001** | **1.25(1.09-1.44)** | **0.001** |
|  | Peak SVG elevation | **1.13(1.02-1.26)** | **0.018** | 1.11(0.95-1.30) | 0.173 |
|  | Area SVG elevation | **1.16(1.06-1.28)** | **0.002** | 1.12(0.96-1.30) | 0.156 |
|  | Peak SVG azimuth | **1.14(1.05-1.23)** | **0.002** | **1.19(1.03-1.38)** | **0.019** |
|  | Area SVG azimuth | 1.04(0.95-1.14) | 0.404 | 1.13(0.96-1.33) | 0.153 |
|  | Peak SVG magnitude | 0.90(0.81-1.01) | 0.084 | 1.00(0.86-1.16) | 0.948 |
|  | Area SVG magnitude | 0.97(0.87-1.08) | 0.580 | 1.10(0.94-1.28) | 0.229 |
|  | SAI QRST | 1.07(0.97-1.17) | 0.168<sup>#</sup> | **1.32(1.17-1.49)** | **<0.0001** |
|  | Heart rate | 1.11(1.00-1.24) | 0.057 | **1.25(1.10-1.43)** | **0.001** |
|  | Bazett's QTc | **1.13(1.01-1.27)** | **0.034** | **1.18(1.09-1.26)** | **<0.0001** |
|  | QRS duration | 0.99(0.89-1.10) | 0.843 | **1.24(1.08-1.42)** | **0.002** |
|  | Cornell voltage | **1.13(1.03-1.25)** | **0.014** | **1.22(1.05-1.42)** | **0.009** |
| Model 2 | Peak QRS-T angle | **1.26(1.15-1.37)** | **<0.0001<sup>#</sup>** | **1.27(1.12-1.44)** | **<0.0001** |
|  | Area QRS-T angle | **1.28(1.16-1.41)** | **<0.0001<sup>#</sup>** | **1.30(1.15-1.48)** | **<0.0001** |
|  | Peak SVG elevation | **1.15(1.04-1.26)** | **0.005** | 1.09(0.95-1.26) | 0.215 |
|  | Area SVG elevation | **1.11(1.01-1.23)** | **0.026** | 1.06(0.92-1.23) | 0.426 |
|  | Peak SVG azimuth | **1.17(1.08-1.26)** | **<0.0001<sup>#</sup>** | **1.26(1.10-1.46)** | **0.001** |
|  | Area SVG azimuth | **1.11(1.01-1.22)** | **0.024<sup>#</sup>** | **1.28(1.09-1.50)** | **0.002** |
|  | Peak SVG magnitude | 0.93(0.83-1.04) | 0.193 | 1.07(0.93-1.23) | 0.355 |
|  | Area SVG magnitude | 0.97(0.87-1.08) | 0.590 | 1.14(0.98-1.31) | 0.080 |
|  | SAI QRST | **1.09(1.03-1.16)** | **0.005** | **1.19(1.07-1.32)** | **0.001** |
|  | Heart rate | 1.11(1.00-1.23) | 0.057 | **1.26(1.11-1.41)** | **<0.0001** |
|  | Bazett's QTc | **1.17(1.08-1.27)** | **<0.0001** | **1.19(1.08-1.31)** | **0.001** |
|  | QRS duration | **1.21(1.12-1.31)** | **<0.0001** | **1.16(1.04-1.30)** | **0.010** |
|  | Cornell voltage | **1.10(1.02-1.18)** | **0.012<sup>#</sup>** | **1.24(1.13-1.37)** | **<0.0001** |
| Model 3 | Peak QRS-T angle | **1.20(1.09-1.33)** | **<0.0001** | **1.17(1.02-1.34)** | **0.028** |
|  | Area QRS-T angle | **1.22(1.09-1.37)** | **<0.0001** | **1.18(1.02-1.37)** | **0.025** |
|  | Peak SVG elevation | 1.08(0.98-1.20) | 0.136 | 0.96(0.82-1.12) | 0.595 |
|  | Area SVG elevation | 1.06(0.96-1.17) | 0.240 | 0.98(0.84-1.15) | 0.823 |
|  | Peak SVG azimuth | **1.13(1.04-1.23)** | **0.005** | **1.16(0.99-1.36)** | **0.062** |
|  | Area SVG azimuth | **1.10(1.003-1.21)** | **0.042** | **1.19(1.02-1.40)** | **0.030** |
|  | Peak SVG magnitude | 1.02(0.94-1.12) | 0.599 | **1.16(1.001-1.35)** | **0.049** |
|  | Area SVG magnitude | 0.95(0.84-1.06) | 0.354 | 1.04(0.91-1.20) | 0.544 |
|  | SAI QRST | 0.97(0.87-1.09) | 0.646 | 1.13(0.98-1.30) | 0.100 |

**Supplemental Table 3A.** Sex interaction in association of GEH with SCD and nonSCD in competing risk models

| Predictor, per 1 SD |  | SCD (n=14,725; 530 SCDs) |  |  |  | nonSCD (n=14,725; 2,178 nonSCDs) |  |  |  |
| --- | --- | --- | --- | --- | --- | --- | --- | --- | --- |
|  |  | SHR(95%CI) | P-value | RSHR for women vs.men (95% CI) | P<sub>interaction</sub> | SHR(95%CI) | P-value | RSHR for women vs.men (95% CI) | P<sub>interaction</sub> |
| Model 1 | Peak QRS-T angle | **1.17(1.07-1.28)** | **<0.0001** | **1.12(0.98-1.28)** | **0.084** | **1.12(1.06-1.19)** | **<0.0001** | 1.12(0.97-1.14) | 0.207 |
|  | Area QRS-T angle | **1.19(1.08-1.31)** | **<0.0001** | 0.98(0.85-1.14) | 0.840 | **1.15(1.08-1.22)** | **<0.0001** | 1.07(0.98-1.18) | 0.120 |
|  | Peak SVG elevation | 1.10(0.99-1.22) | 0.084 | 1.07(0.90-1.28) | 0.445 | 1.03(0.97-1.10) | 0.339 | 0.97(0.88-1.06) | 0.474 |
|  | Area SVG elevation | **1.13(1.03-1.23)** | **0.011** | 1.06(0.90-1.26) | 0.484 | 1.01(0.95-1.07) | 0.752 | 0.99(0.91-1.08) | 0.784 |
|  | Peak SVG azimuth | **1.10(1.02-1.20)** | **0.020** | 1.07(0.92-1.24) | 0.408 | 1.04(0.99-1.10) | 0.115 | 1.06(0.98-1.16) | 0.144 |
|  | Area SVG azimuth | 1.01(0.92-1.10) | 0.884 | 1.07(0.86-1.32) | 0.557 | **1.07(1.01-1.14)** | **0.022** | 1.03(0.94-1.13) | 0.533 |
|  | Peak SVG magnitude | **0.89(0.79-0.99)** | **0.040** | **1.16(0.98-1.36)** | **0.083** | **0.92(0.86-0.99)** | **0.019** | **1.11(1.01-1.22)** | **0.027** |
|  | Area SVG magnitude | 0.94(0.84-1.05) | 0.250 | **1.16(0.99-1.37)** | **0.063** | 0.94(0.88-1.00) | 0.061 | **1.15(1.05-1.26)** | **0.003** |
|  | SAI QRST | 1.03(0.93-1.13) | 0.604 | **1.16(1.01-1.33)** | **0.033** | 0.99(0.93-1.06) | 0.728 | **1.16(1.06-1.27)** | **0.002** |
|  | Heart rate. | 1.01(0.91-1.12) | 0.914 | **1.16(0.97-1.37)** | **0.087** | **1.18(1.11-1.26)** | **<0.0001** | 0.98(0.89-1.07) | 0.624 |
|  | Bazett's QTc | 1.02(0.92-1.14) | 0.697 | **1.15(1.02-1.30)** | **0.025** | **1.20(1.12-1.28)** | **<0.0001** | **0.87(0.81-0.94)** | **0.001** |
|  | QRS duration | 0.98(0.89-1.08) | 0.695 | **1.27(1.08-1.49)** | **0.004** | 1.05(0.99-1.12) | 0.079 | 0.95(0.88-1.03) | 0.197 |
|  | Cornell voltage | 1.06(0.96-1.16) | 0.249 | 1.14(0.97-1.34) | 0.112 | **1.10(1.03-1.17)** | **0.003** | **1.11(1.01-1.22)** | **0.032** |
| Model 2 | Peak QRS-T angle | **1.10(1.01-1.21)** | **0.039** | **1.13(0.98-1.31)** | **0.081** | **0.90(0.85-0.96)** | **0.001** | **1.10(1.004-1.20)** | **0.040** |
|  | Area QRS-T angle | **1.15(1.03-1.27)** | **0.007** | 1.09(0.93-1.27) | 0.296 | **0.90(0.85-0.96)** | **0.001** | 1.09(0.99-1.20) | 0.078 |
|  | Peak SVG elevation | 1.10(1.00-1.22) | 0.061 | 1.06(0.89-1.27) | 0.480 | 0.94(0.88-1.01) | 0.082 | 0.99(0.89-1.10) | 0.833 |
|  | Area SVG elevation | 1.07(0.97-1.18) | 0.189 | 1.10(0.93-1.30) | 0.283 | 0.97(0.90-1.04) | 0.353 | 0.95(0.86-1.06) | 0.366 |
|  | Peak SVG azimuth | **1.09(1.002-1.19)** | **0.044** | **1.16(0.99-1.35)** | **0.065** | **0.94(0.89-0.99)** | **0.026** | 1.05(0.95-1.15) | 0.379 |
|  | Area SVG azimuth | 1.03(0.94-1.14) | 0.506 | **1.21(0.99-1.48)** | **0.063** | 0.95(0.90-1.01) | 0.083 | 1.02(0.93-1.12) | 0.605 |
|  | Peak SVG magnitude | 0.94(0.84-1.05) | 0.277 | **1.18(1.00-1.39)** | **0.057** | 1.02(0.95-1.10) | 0.605 | 0.97(0.87-1.07) | 0.524 |
|  | Area SVG magnitude | 0.95(0.85-1.06) | 0.341 | **1.21(1.03-1.43)** | **0.021** | 1.00(0.93-1.08) | 0.904 | 1.00(0.90-1.10) | 0.933 |
|  | SAI QRST | 1.05(0.97-1.13) | 0.205 | **1.16(1.02-1.31)** | **0.025** | **0.94(0.89-0.99)** | **0.019** | 1.01(0.93-1.10) | 0.810 |
|  | Heart rate. | 0.98(0.88-1.09) | 0.714 | **1.19(1.01-1.40)** | **0.036** | **1.08(1.01-1.15)** | **0.032** | 1.06(0.96-1.17) | 0.230 |
|  | Bazett's QTc | 1.07(0.98-1.17) | 0.124 | **1.11(0.99-1.26)** | **0.086** | 1.01(0.95-1.07) | 0.714 | 1.02(0.94-1.12) | 0.619 |
|  | QRS duration | 1.13(1.04-1.23) | 0.002 | 1.10(0.96-1.27) | 0.176 | **0.94(0.89-0.99)** | **0.032** | 0.94(0.85-1.03) | 0.154 |
|  | Cornell voltage | 1.03(0.95-1.11) | 0.442 | **1.23(1.10-1.38)** | **<0.0001** | 0.97(0.93-1.02) | 0.271 | 0.97(0.90-1.06) | 0.525 |

RSHR=relative sub-hazard ratio

**Supplemental Table 3B.**
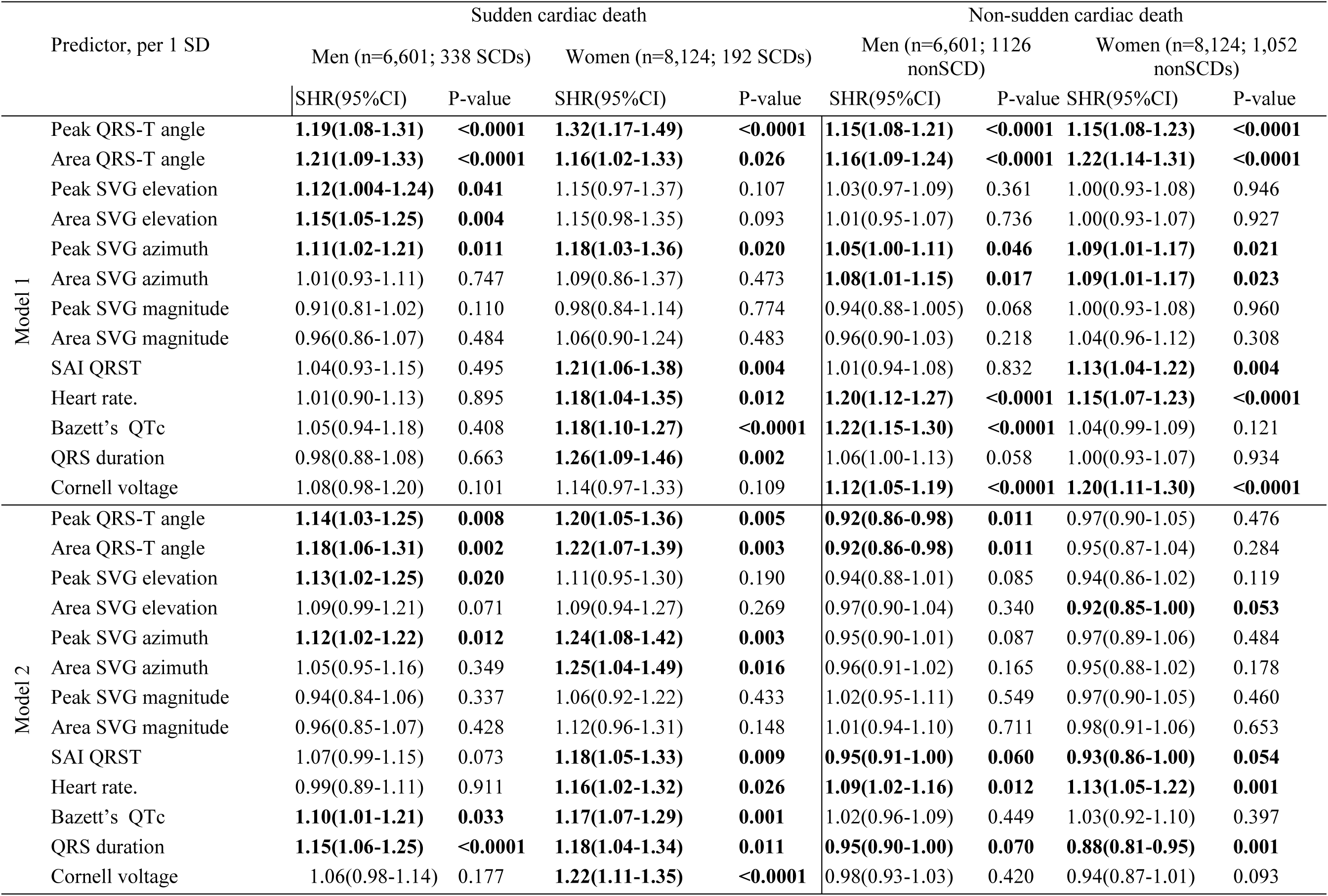
Competing risks of sudden cardiac death and non-sudden cardiovascular death for men and women

### Supplemental Figure Legends

**Supplemental Figure 1.** Estimated adjusted marginal (least-squares) means and 95% Confidence Intervals of GEH variables for men and women. **Model 1** was adjusted for age, race, and study center. **Model 2** was in addition adjusted for prevalent HF, CHD, stroke, diabetes, hypertension, BMI, postmenopause state, education level, current smoking, current alcohol intake, leisure physical activity level, BMI category, use of antihypertensive and antiarrhythmic medications, levels of total cholesterol, HDL, and triglycerides, serum concentrations of sodium, potassium, calcium, magnesium, phosphorus, and uric acid, total protein and albumin, blood urea nitrogen, CKD stage classified by eGFRCKD-EPI, mean heart rate, QRS duration, QTc, Cornell voltage, and sex-specific ECG – LVH.

**Supplemental Figure 2.** Adjusted (for age, race, study center, prevalent at baseline HF, CHD, stroke, diabetes, hypertension, BMI, postmenopause state, education level, current smoking, current alcohol intake, leisure physical activity level, BMI category, use of antihypertensive and antiarrhythmic medications, levels of total cholesterol, HDL, and triglycerides, serum concentrations of sodium, potassium, calcium, magnesium, phosphorus, and uric acid, total protein and albumin, blood urea nitrogen, CKD stage classified by eGFRCKD-EPI, mean heart rate, QRS duration, QTc, Cornell voltage, and sex-specific ECG – LVH.) risk of SCD associated with GEH variables in men and women. Restricted cubic spline with 95% CI shows change in hazard ratio (Y-axis) in response to GEH variable change (X-axis). 50^th^ percentile of GEH variable is selected as reference.

**Figure 1A:**
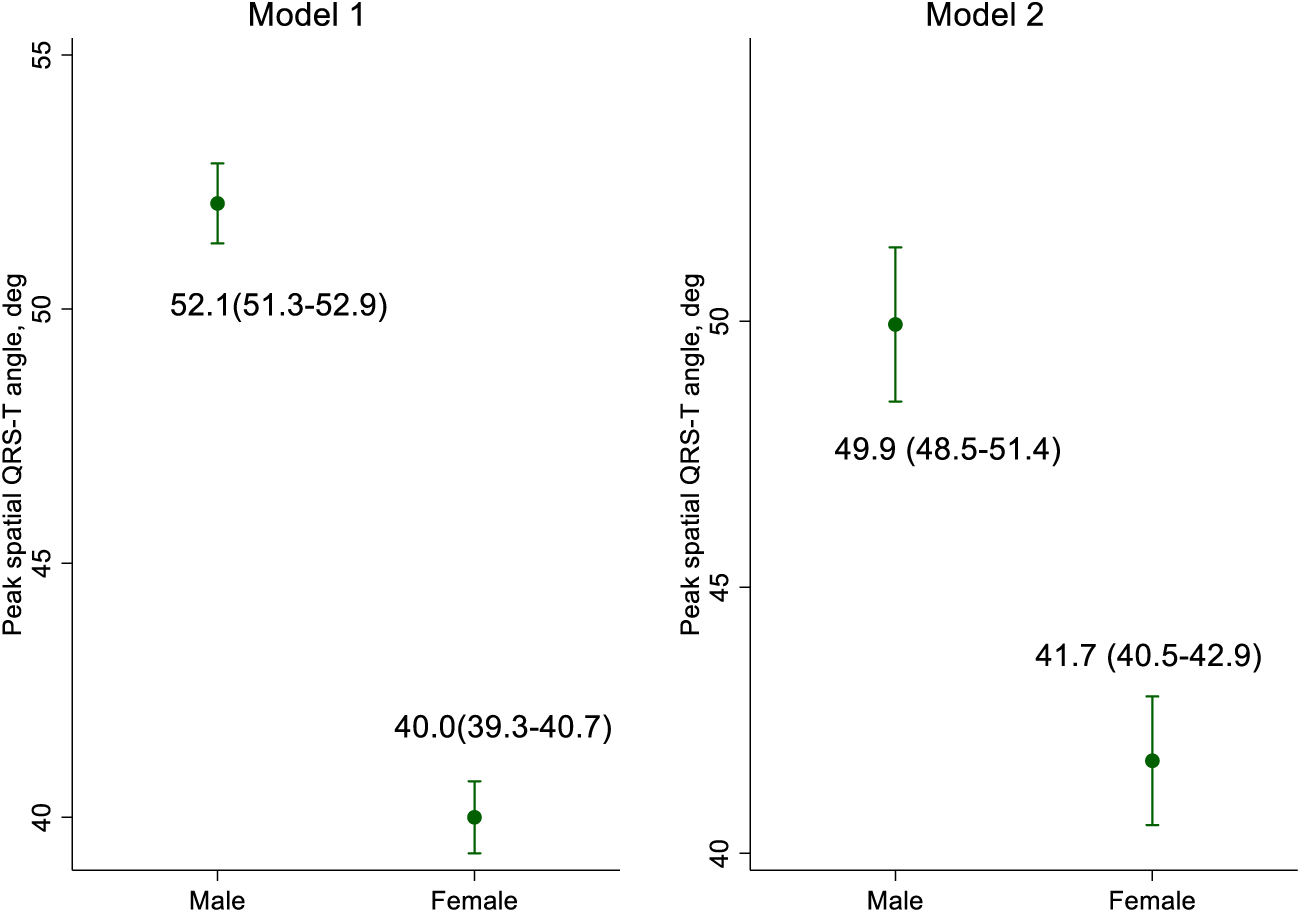
Spatial peak QRS-T angle in men and women:

**Figure 1B:**
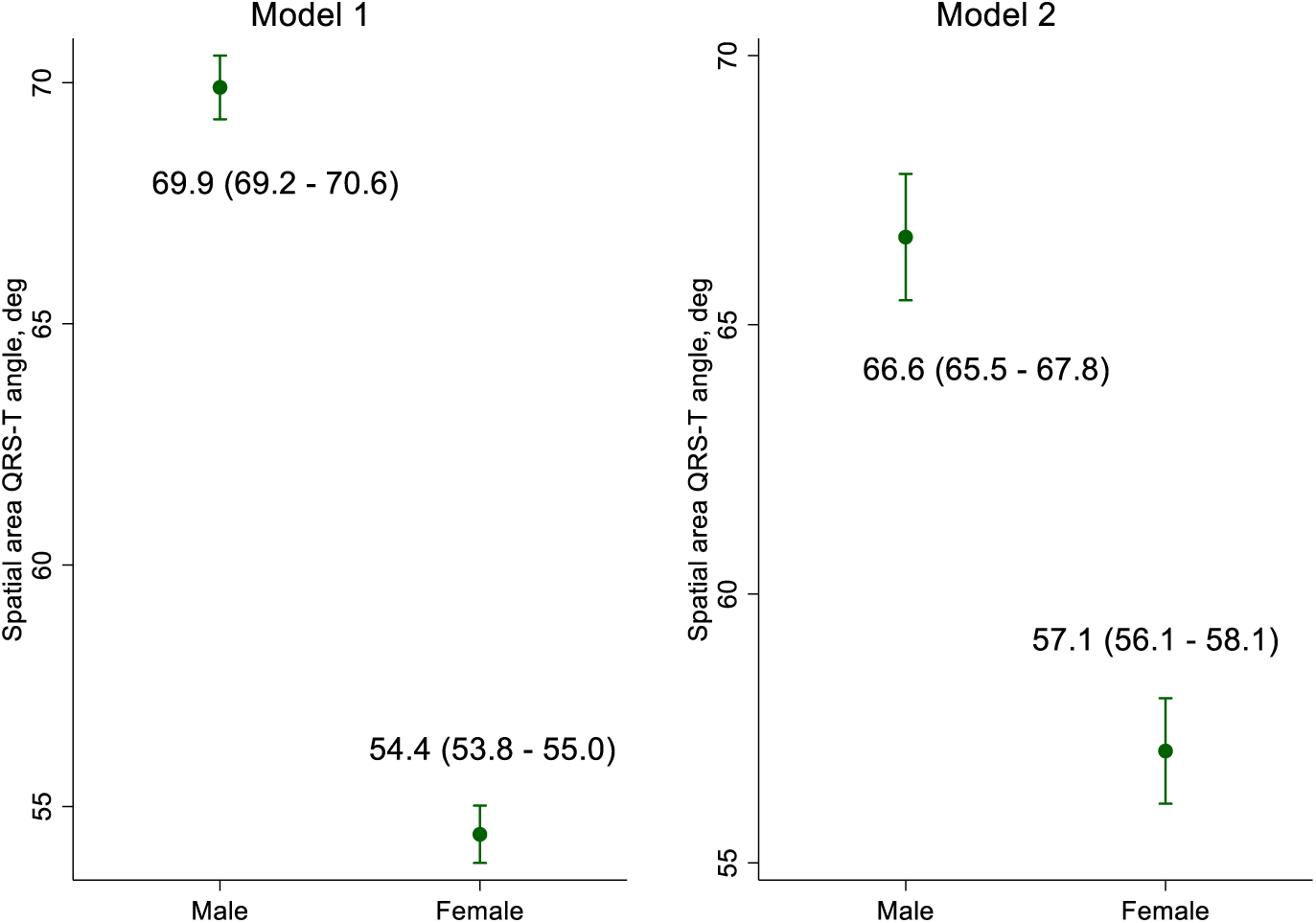
Spatial area QRS-T angle in men and women:

**Figure 1C:**
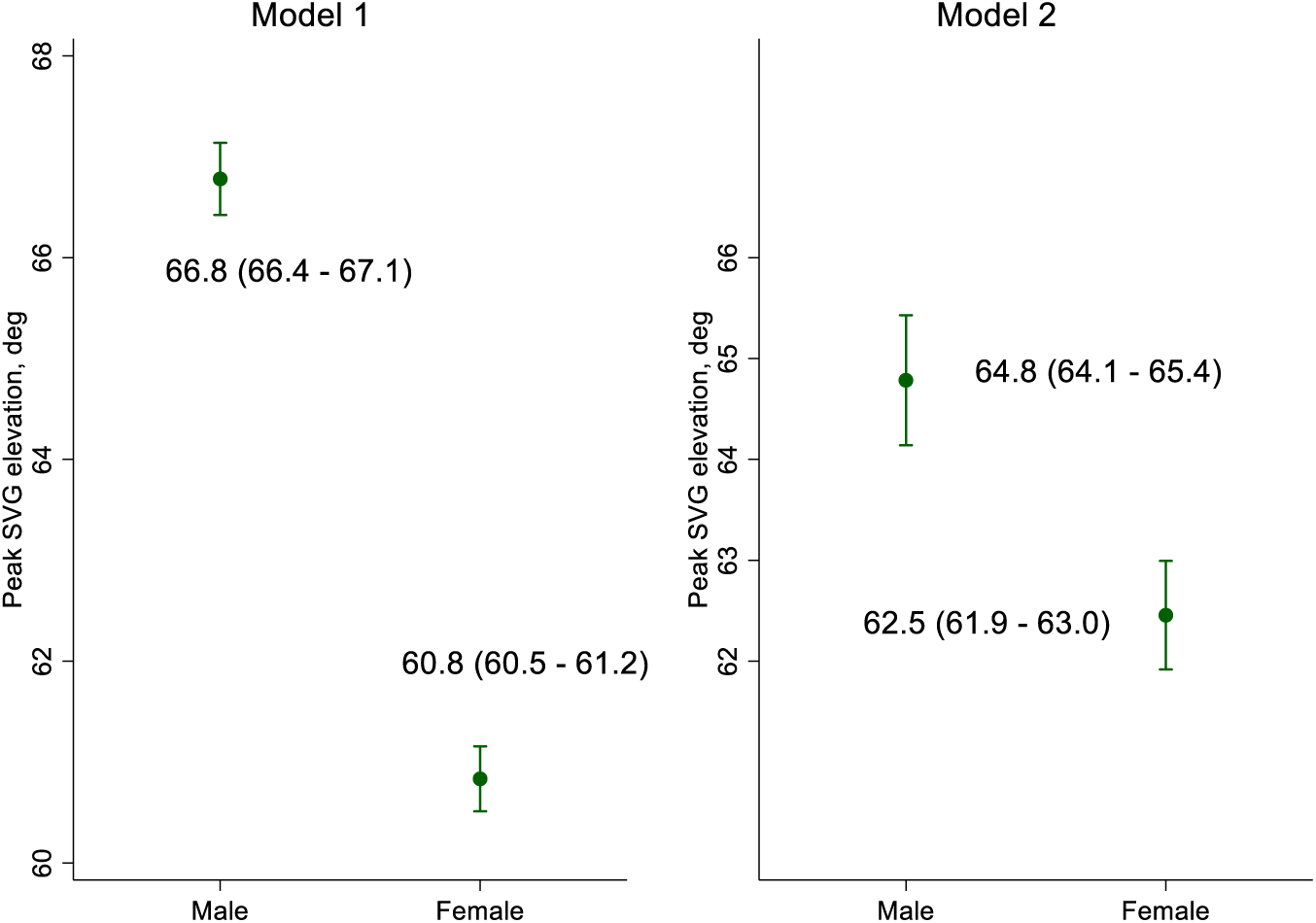
Spatial peak SVG elevation in men and women:

**Figure 1D:**
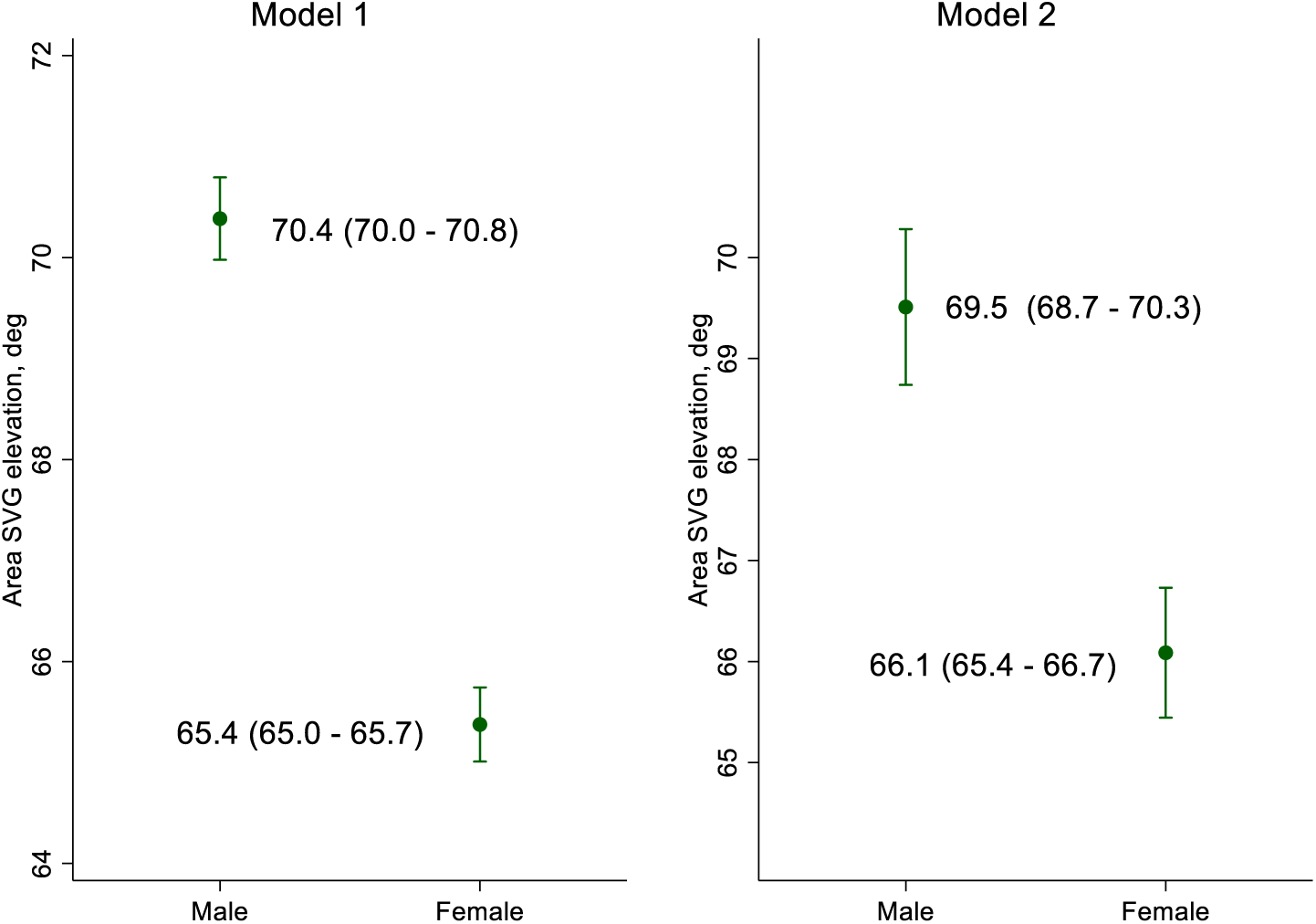
Spatial area SVG elevation in men and women:

**Figure 1E:**
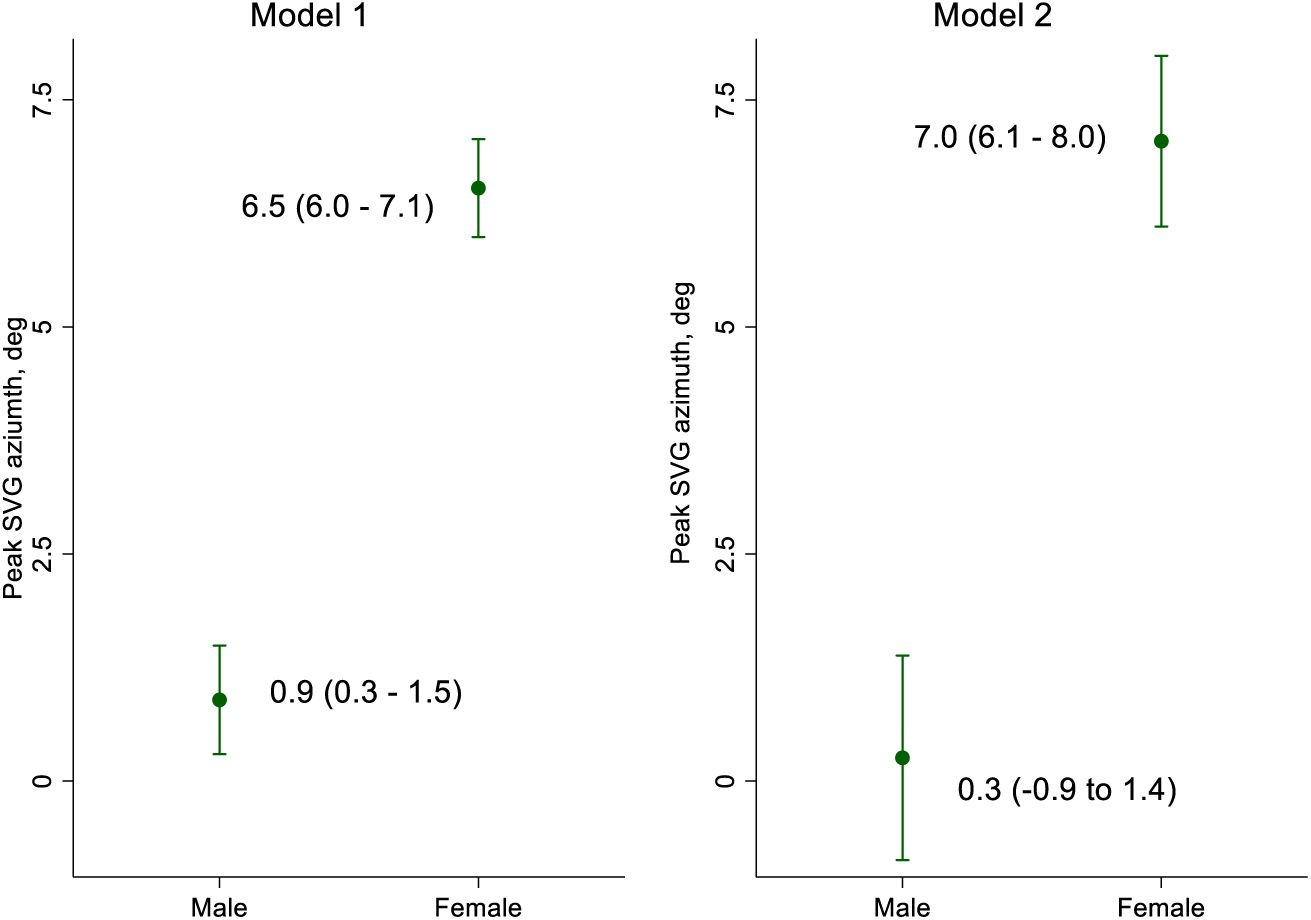
Spatial peak SVG azimuth in men and women:

**Figure 1F:**
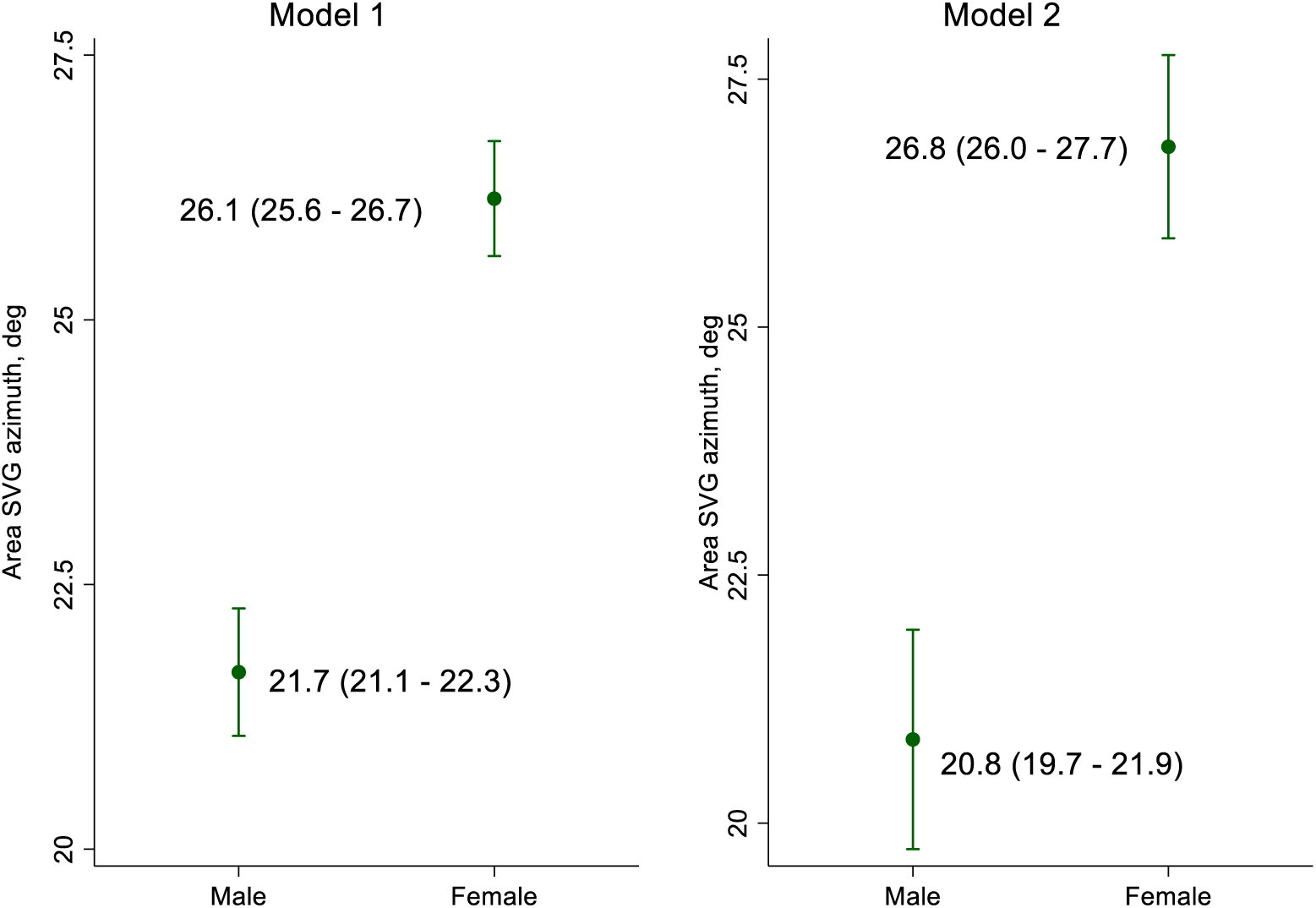
Spatial area SVG azimuth in men and women:

**Figure 1G:**
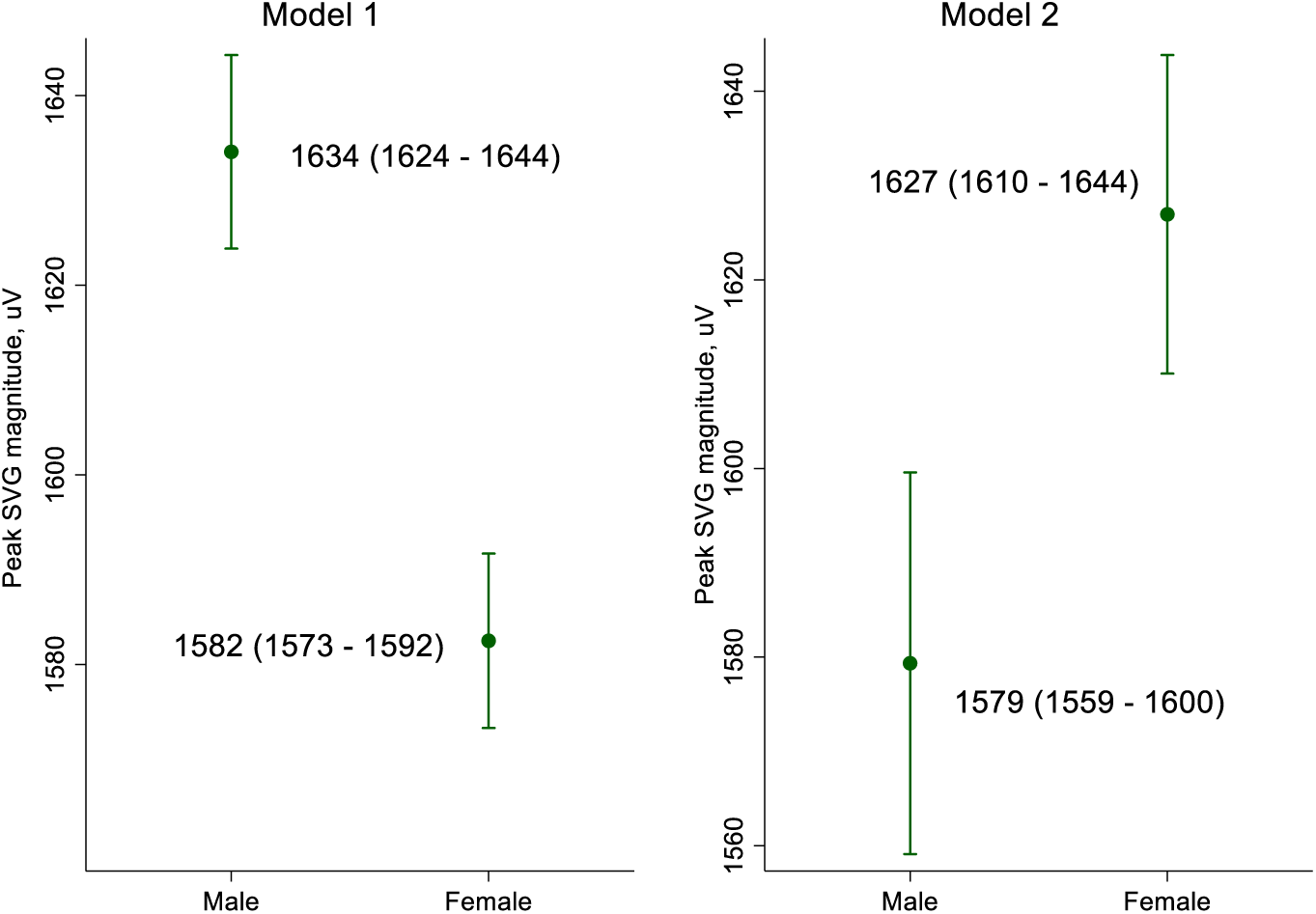
Spatial peak SVG magnitude in men and women:

**Figure 1H:**
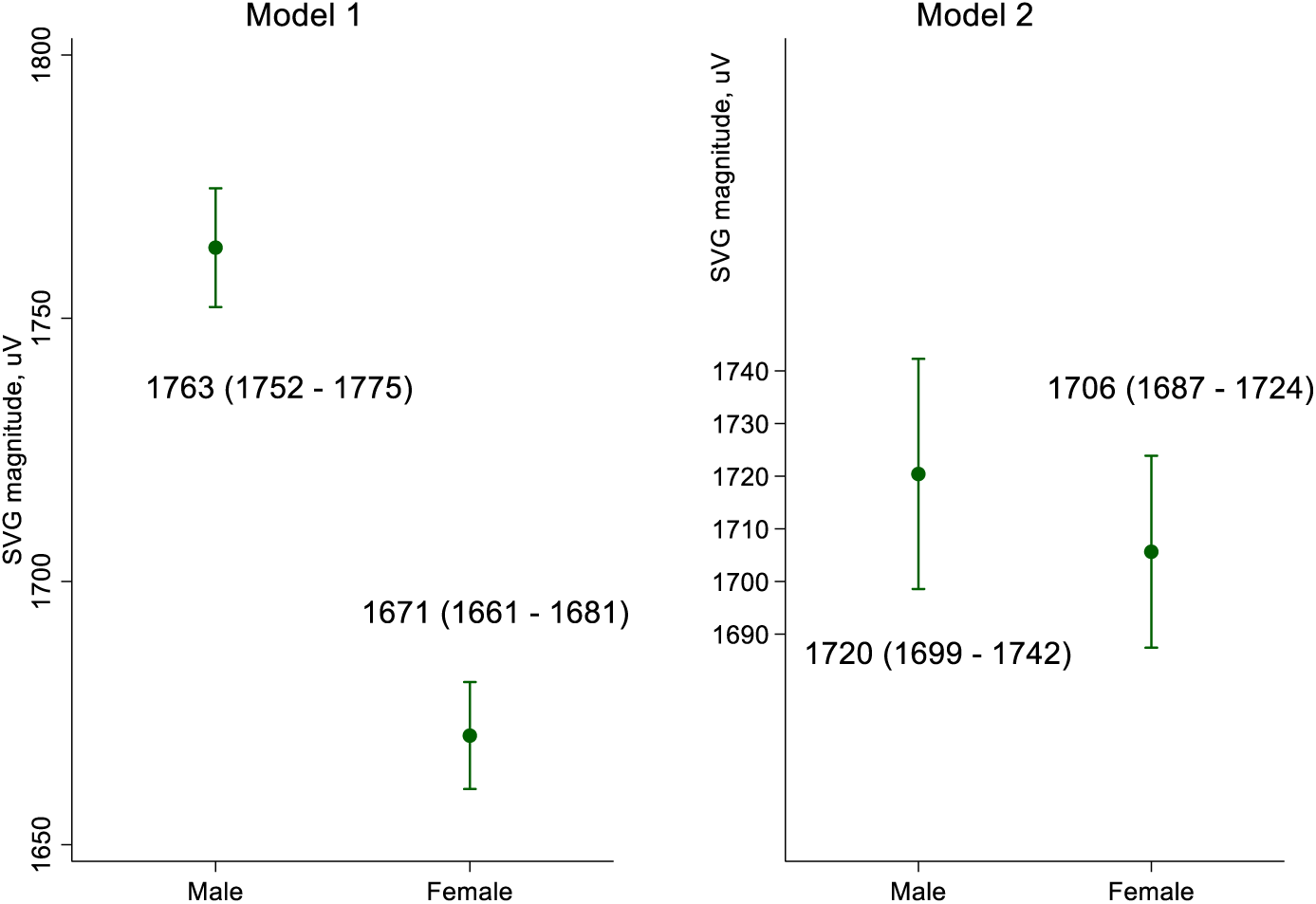
Spatial SVG magnitude in men and women:

**Figure 1I:**
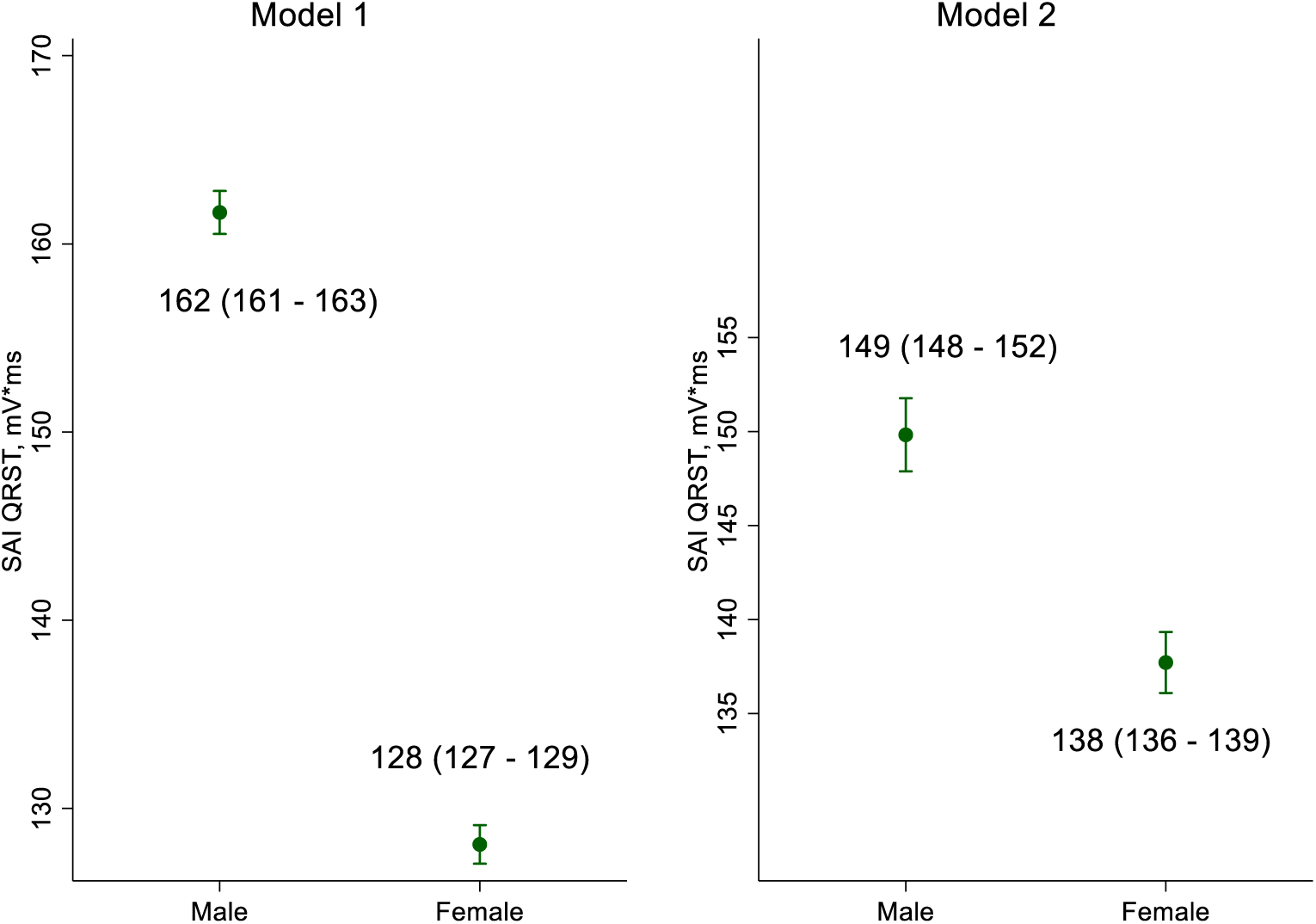
SAI QRST in men and women:

**Figure 2A:**
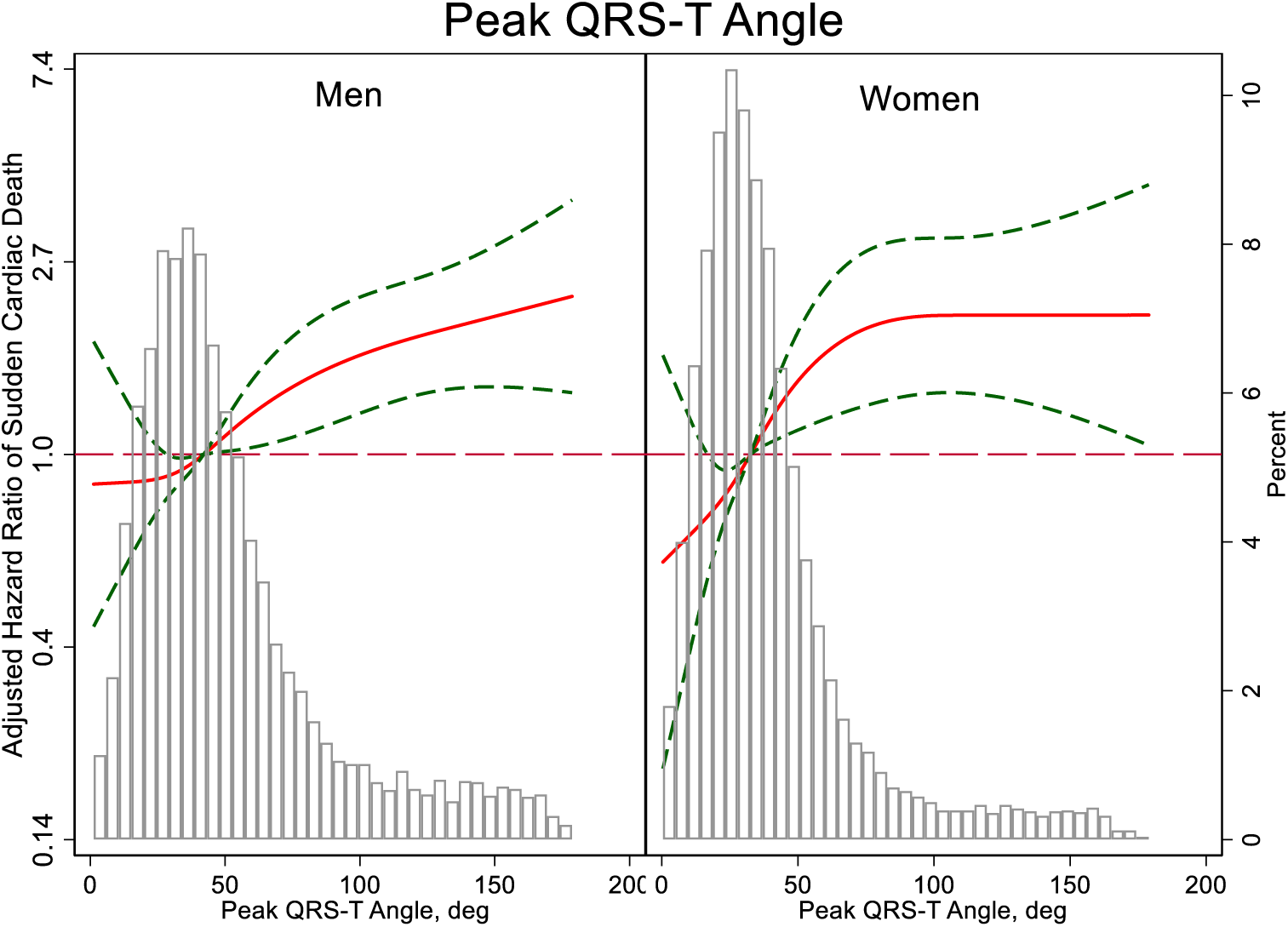
Adjusted risk of SCD associated with peak QRS-T angle

**Figure 2B:**
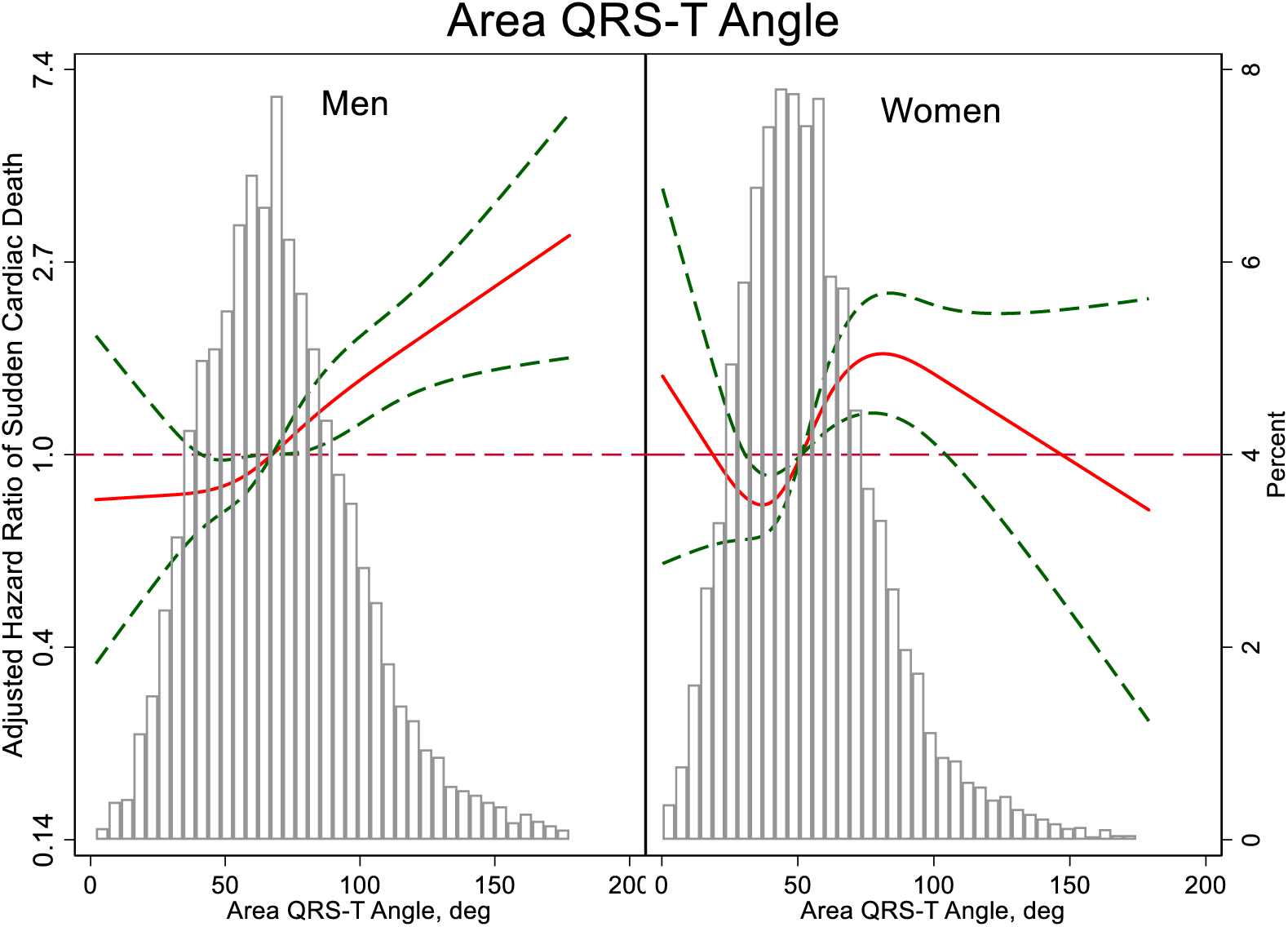
Adjusted risk of SCD associated with area QRS-T angle

**Figure 2C:**
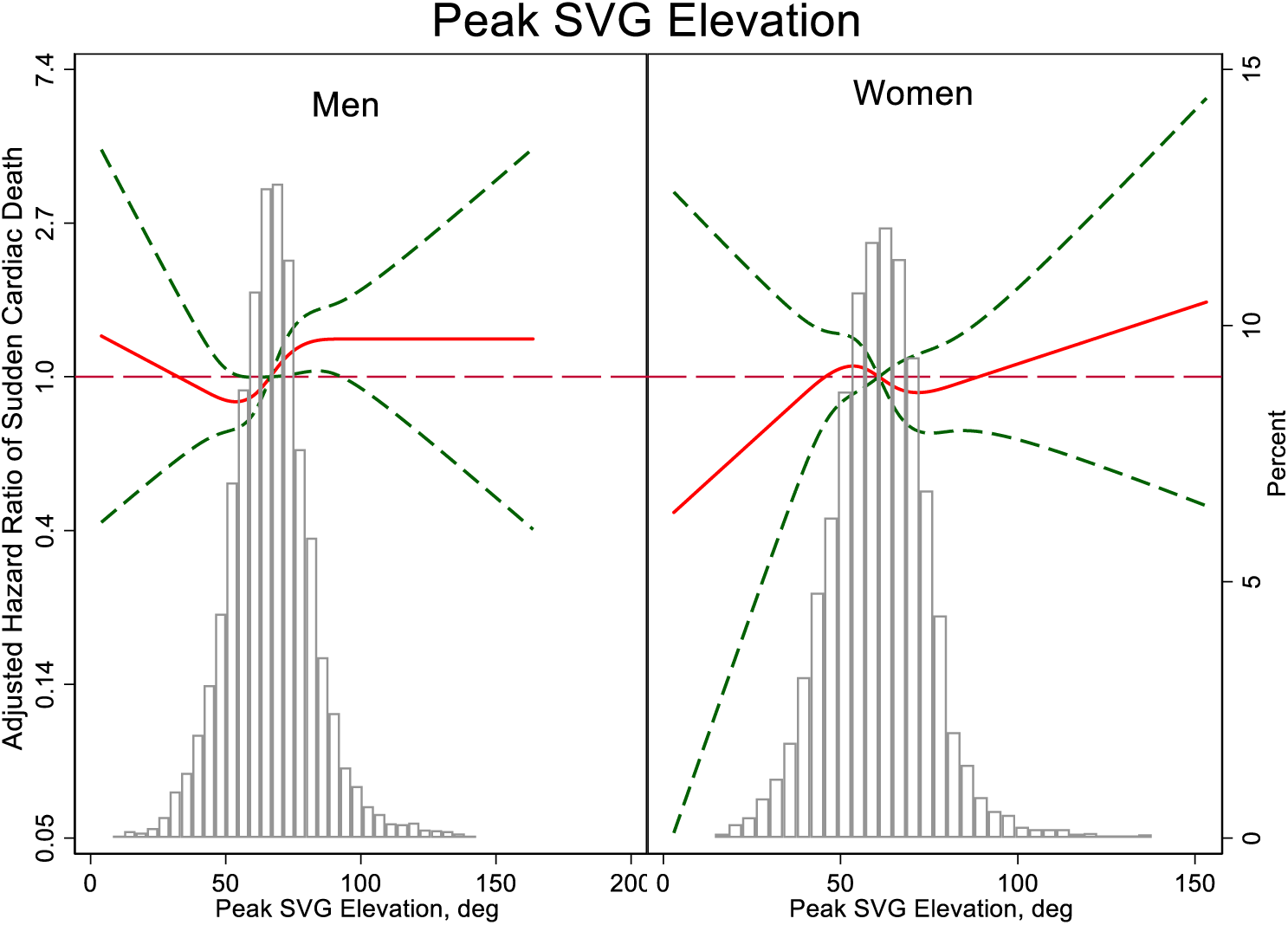
Adjusted risk of SCD associated with peak SVG elevation

**Figure 2D:**
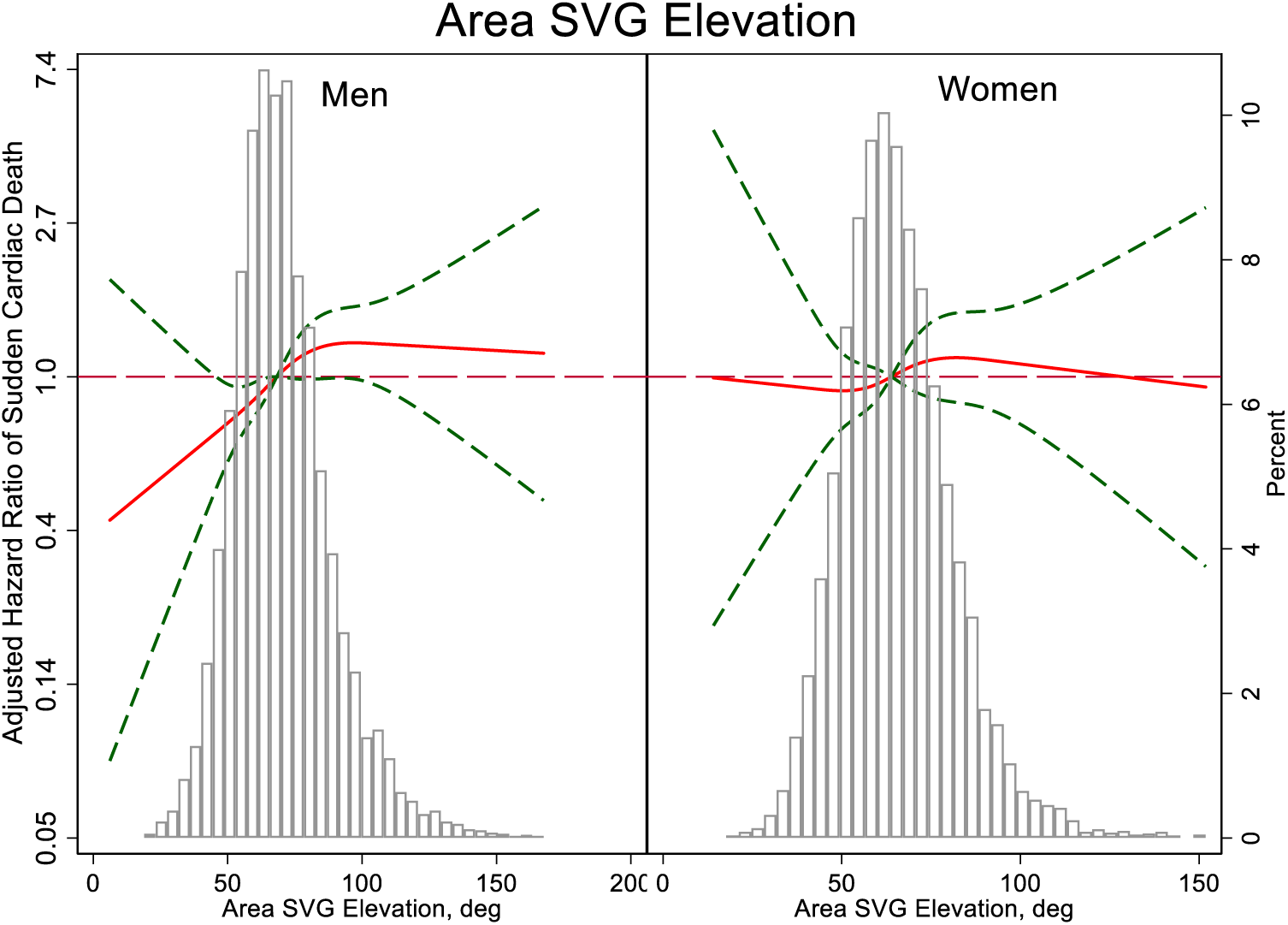
Adjusted risk of SCD associated with area SVG elevation

**Figure 2E:**
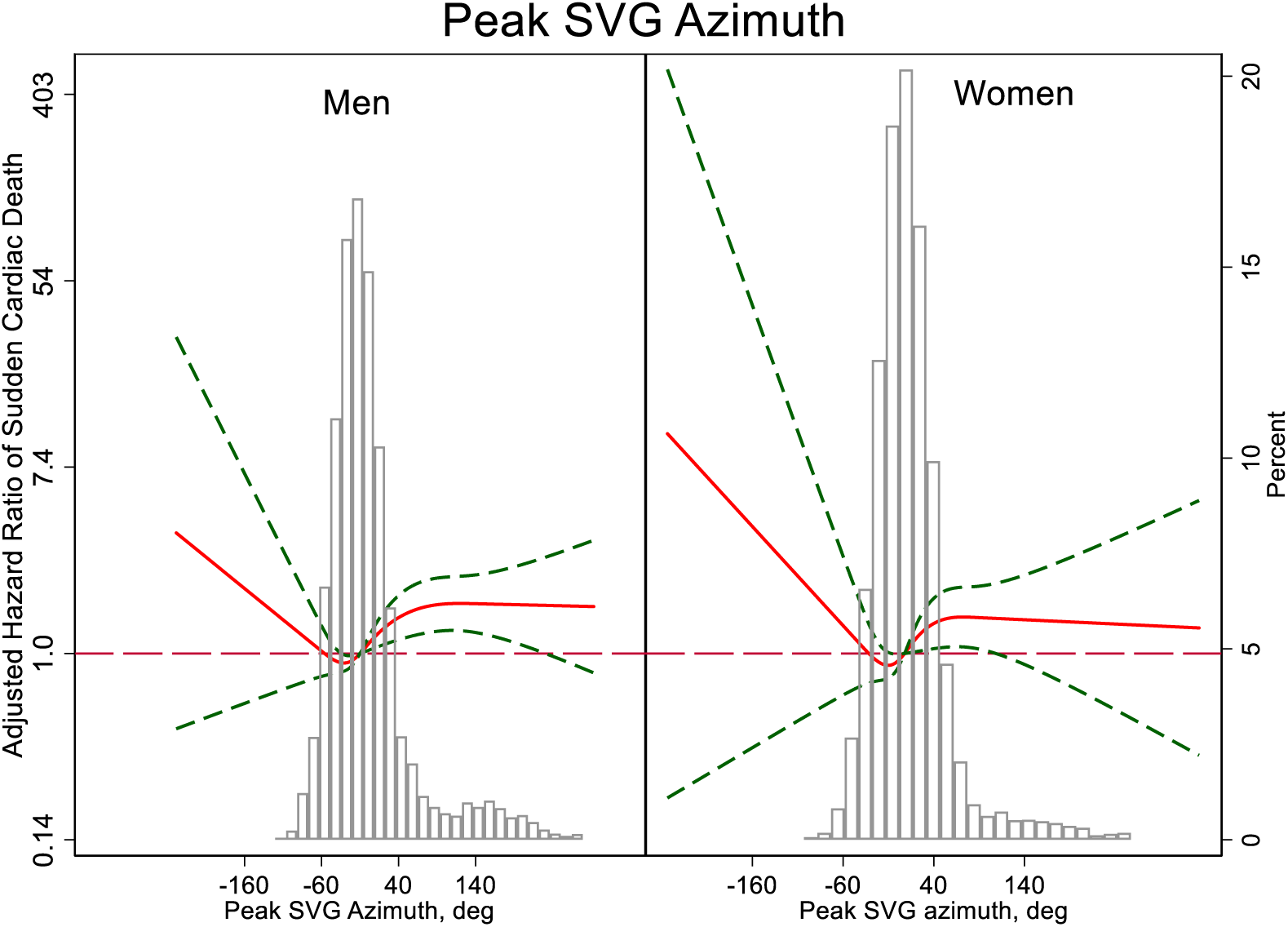
Adjusted risk of SCD associated with peak SVG azimuth

**Figure 2F:**
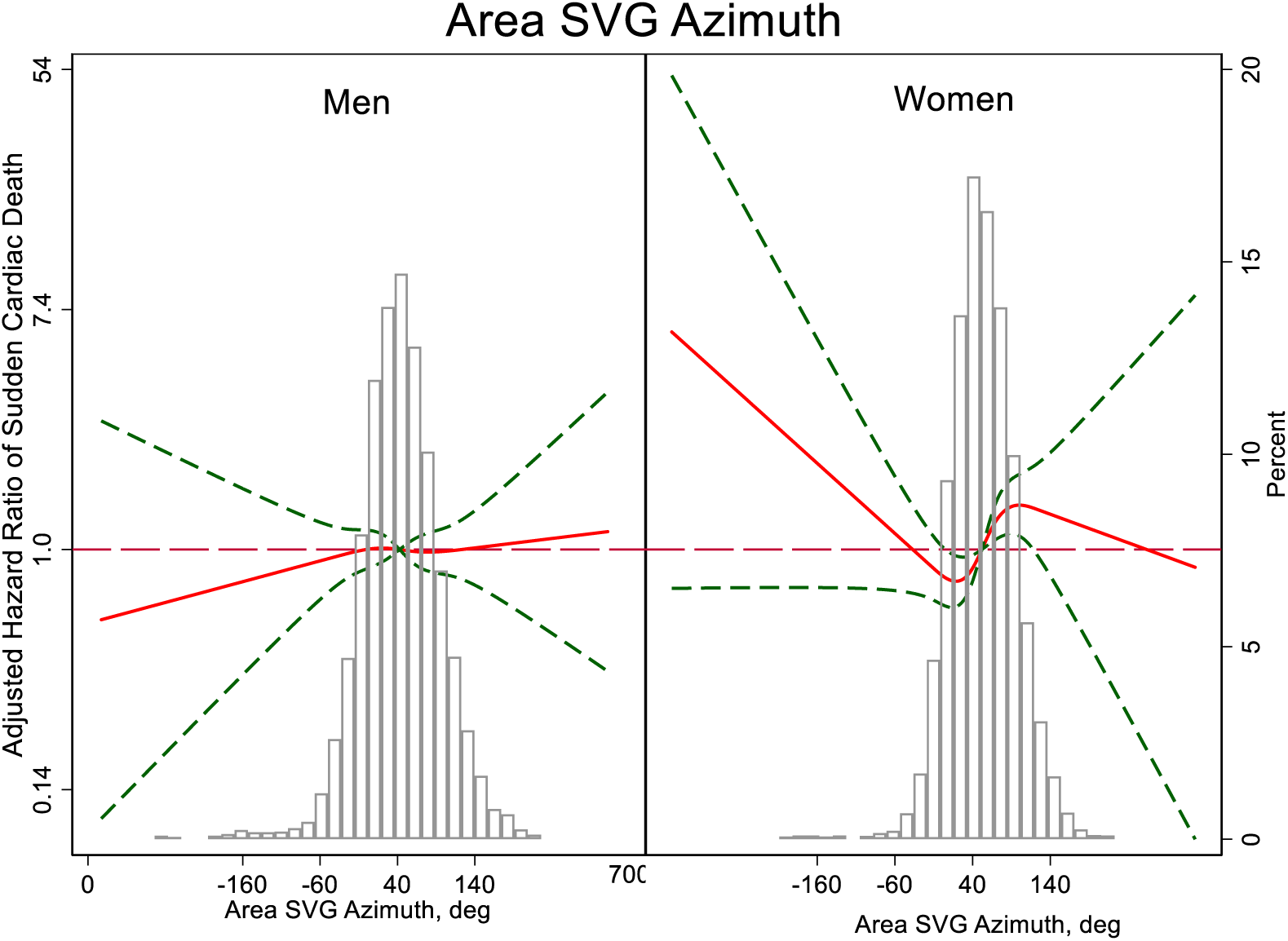
Adjusted risk of SCD associated with area SVG azimuth

**Figure 2G:**
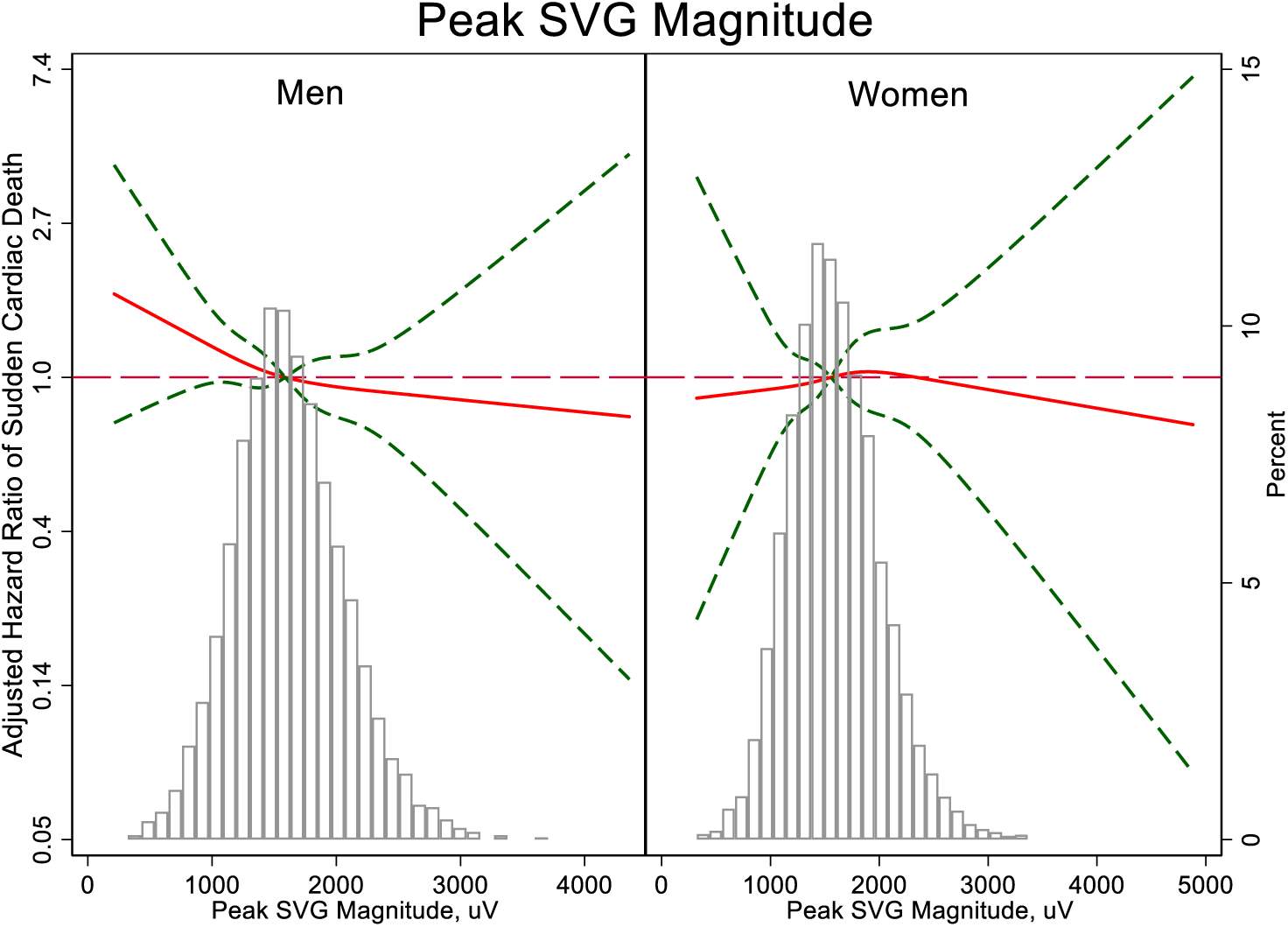
Adjusted risk of SCD associated with peak SVG magnitude

**Figure 2H:**
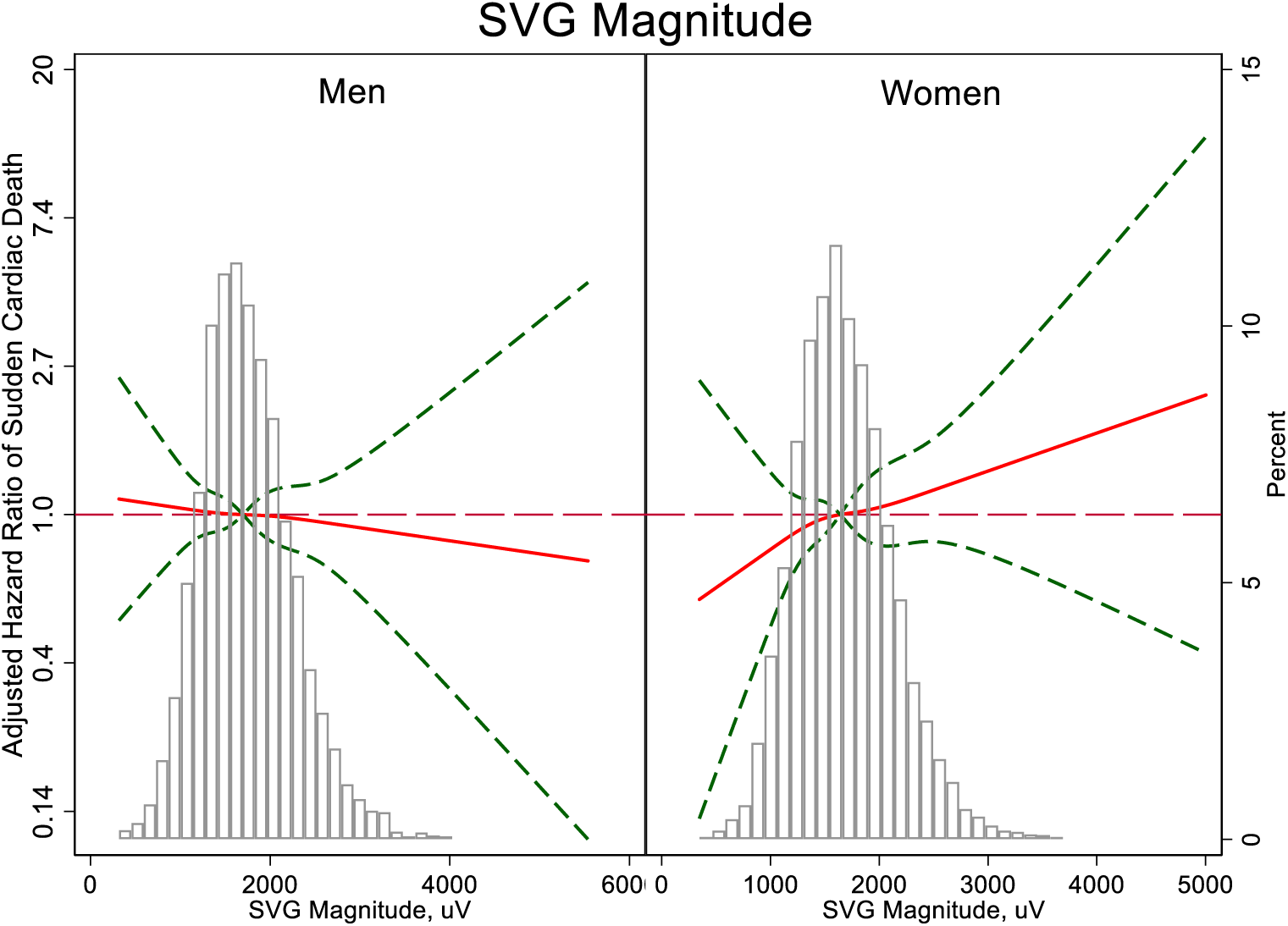
Adjusted risk of SCD associated with SVG magnitude

**Figure 2I:**
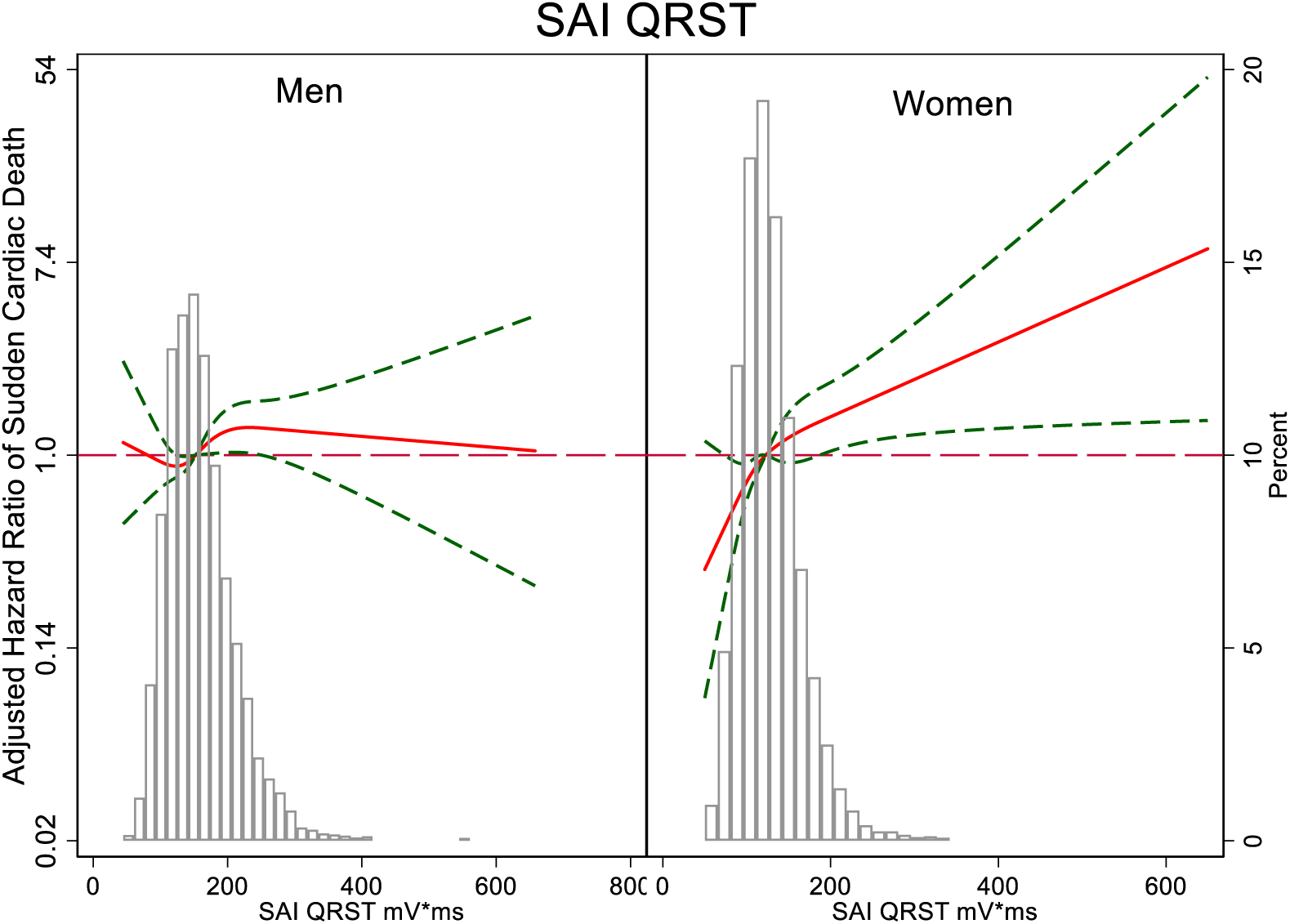
Adjusted risk of SCD associated with SAI QRST

